# Retroelement co-option disrupts the cancer transcriptional programme

**DOI:** 10.1101/2025.02.21.639580

**Authors:** Jane Loong, Rachael Thompson, Callum Hall, Laura Doglio, Judith Pape, George Kassiotis

**Author notes:** These authors contributed equally to this work (Jane Loong, Rachael Thompson). Correspondence: Dr George Kassiotis.

## Abstract

**Background:** Transcriptional activation of otherwise repressed endogenous retroelements (RTEs) is a hallmark of cancer, shaping tumour progression and immunogenicity by multifaceted, yet incompletely understood mechanisms.

**Methods:** We used an extended pan-cancer transcriptome assembly to identify potential effects of RTEs on the genes near or within which they have integrated. These were subsequently verified in test cases by further analysis of transcriptional profiles in cancer patient data, and by *in vitro* studies involving restoration of gene activity, and proliferation and migration assays in cancer cell lines.

**Results:** We report that cancer-specific transcriptional activation of RTEs causes frequent reduction or loss of gene function. Exonisation and alternative splicing of RTEs creates non-functional RNA and protein isoforms and derepressed RTE promoter activity initiates antisense transcription, both at the expense of the canonical isoforms. Contrary to theoretical expectation, transcriptionally activated RTEs affect genes with established tumour-promoting function, including the common essential *RNGTT* and the lung cancer-promoting *CHRNA5* genes. Furthermore, the disruptive effect of RTE activation on adjacent tumour-promoting genes is associated with slower disease progression in clinical data, whereas experimental restoration of gene activity enhances tumour cell *in vitro* growth and invasiveness

**Conclusions:** These findings underscore the gene-disruptive potential of seemingly innocuous germline RTE integrations, unleashed only by their transcriptional utilisation in cancer. They further suggest that such metastable RTE integrations are co-opted as sensors of the epigenetic and transcriptional changes occurring during cellular transformation and as executors that disrupt the function of tumour-promoting genes.

## Introduction

Similar to all mammalian genomes, the human genome has amassed over 4 million recognisable integrations of retrotransposable elements (RTEs) of diverse subfamilies, some of which are primate-specific, such as the human endogenous retrovirus H (*HERVH*) subfamily of long terminal repeat (LTR) RTEs and the *Alu* subfamily of non-LTR RTEs [1]. While the vast majority of human RTEs have lost the ability to retrotranspose, new germline integrations of active RTEs are acquired slowly but continually [2].

RTE integration poses a significant risk of insertional mutagenesis, which is higher for the recently-acquired or active subfamilies. Highly deleterious mutations may be quickly counterselected, whereas less damaging integrations are retained for longer evolutionary times. The genetic diversity generated by RTE integration, as well as the regulatory sequences they carry, can be co-opted in the evolution of new physiological host functions and transcriptional networks [3, 4]. Indeed, RTE enhancer and promoter activities are co-opted in the regulatory networks of placentation or the interferon (IFN) response genes [5], and alternative RTE exons in the diversification of protein isoforms and function [4].

In addition to evolutionary selection, epigenetic control of RTEs further mitigates the potentially damaging effect of RTE integrations on host gene function by preventing their independent transcription or inclusion in host gene transcripts [6]. Epigenetic repression acts faster than evolutionary processes and allows for the latter to determine the ultimate fate of a given RTE integration. However, epigenetic repression is reversible and can be lost, particularly following the major epigenetic changes seen in cancer [6]. In turn, RTE release from epigenetic control may allow for previously hidden effects on the gene near or within which they have integrated to manifest.

Altered transcriptional activity of RTEs has been consistently observed in most cancer types, associated with substantial effects on the cancer transcriptome [6, 7]. RTE transcriptional inclusion in cancer results in aberrant transcription and splicing patterns, often in a cancer type-specific fashion [8]. However, given its high degree of complexity, the functional consequences of the aberrant cancer transcriptome created by RTE derepression is not yet fully understood. RTE can be co-opted in driving elevated or ectopic expression of genes with tumour-promoting, or in creating tumorigenic protein isoforms. Instances of such onco-exaptation events include the elevated expression of *CSF1R* and *IRF5* in Hodgkin’s lymphoma [9, 10], the ectopic expression of *CALB1* in squamous lung cancer [11], and the creation of an oncogenic form of anaplastic lymphoma kinase (ALK) in melanoma [12]. Moreover, comprehensive analysis of epigenetically reactivated RTE promoter activity has identified >100 potential onco-exaptation events involving oncogenes in diverse cancer types [13].

Owing to the increase in tumour cell fitness they confer, RTE-mediated activation of tumour-promoting genes would be increasingly enriched over the course of tumour progression and molecular evolution, making such onco-exaptation events easier to identify in cancer transcriptomes. In contrast, effects of epigenetically reactivated RTEs that compromise the function of an essential gene would also compromise tumour cell fitness and would, therefore, be counterselected during tumour evolution, giving the appearance of rarer occurrence. However, the potential of transcriptionally reactivated RTEs to disrupt the function of tumour-promoting genes may be a selectable trait in host species evolution.

Here, we investigated the degree to which gene function may be compromised by transcriptionally reactivated RTEs in cancer. We made use of our increasing understanding of the complexity of cancer transcriptomes and identified cancer-specific transcript isoforms, created by transcriptional inclusion of RTEs at the expense of the protein-coding canonical isoform of adjacent genes. Counterintuitively, many of the affected genes have an established tumour-promoting role, and, in test cases, restoration of their expression accelerates tumour cell-intrinsic growth and invasiveness.

## Methods

### Transcript identification, read mapping and quantitation

RNA-seq samples were downloaded from The Genotype-Tissue Expression (GTEx) project and TCGA (poly(A) selected RNA), CCLE and other publicly available RNA-seq repositories listed in the Data availability statement. Samples from TCGA were downloaded through the *gdc-client* application. The .bam files were parsed with a custom Bash pipeline using GNU parallel [14], and converted to .fasta files using SAMtools v1.8 [15]. RNA-seq data from TCGA, GTEx, CCLE and listed previous studies were mapped to our *de novo* cancer transcriptome assembly and counted as previously described [8]. Briefly, transcripts per million (TPM) values were calculated for all transcripts in the transcript assembly [8] with a custom Bash pipeline and Salmon v0.8.2 [16], which uses a probabilistic model for assigning reads aligning to multiple transcript isoforms, based on the abundance of reads unique to each isoform [16]. Read count tables were additionally imported into Qlucore Omics Explorer v3.9 (Qlucore, Lund, Sweden) for further downstream expression analyses and visualization. Splice junctions were visualized using the Integrative Genome Viewer (IGV) v2.4.19 [17]. Long-read RNA-seq samples were aligned to the GRCh38/hg38 human genome using minimap2 v2.17 [18]. For assemblies of long-read RNA-seq reads, the obtained .bam files were first converted to bed12 using bam2bed12.py script from FLAIR suit [19]. High-confidence isoforms were selected using “collapse” function from flair.py script [19].

### Hypoxia scores

The hypoxia scores [20] for all TCGA samples were kindly provided by Prof. David Mole (Nuffield Department of Medicine, University of Oxford). The hypoxia scores for KIRC samples were filtered, however mismatches in sample names due to updates by TCGA to the naming system of files meant the mean hypoxia score per patient was used here. Of the 479 KIRC patients with hypoxia scores calculated for tumour samples, 406 had hypoxia scores for one sample, 70 for two samples, and three patients had the mean hypoxia score calculated from three samples.

### Repeat annotation and enrichment analysis

Repeat regions were annotated as previously described [21]. Briefly, hidden Markov models (HMMs) representing known Human repeat families (Dfam 2.0 library v150923) were used to annotate GRCh38 using RepeatMasker, configured with nhmmer. RepeatMasker annotates LTR and internal regions separately, thus tabular outputs were parsed to merge adjacent annotations for the same element. Enrichment analysis of repeat types was performed using Fisher’s Exact test in MATLAB (version R2022b, TheMathWorks), followed by the Bonferroni-Holm method to correct for multiple testing.

### Functional gene annotation by gene ontology

Pathway analyses were performed using g:Profiler (https://biit.cs.ut.ee/gprofiler) with genes ordered by the degree of differential expression. P values were estimated by hypergeometric distribution tests and adjusted by multiple testing correction using the g:SCS (set counts and sizes) algorithm, integral to the g:Profiler server [22].

### Survival analysis and hazard ratio calculations

Overall survival time was downloaded for TCGA data alongside all other metadata. Survival analysis was run using R and RStudio (version 2023.12.0 Build 369). Univariate and multivariate analyses, hazard ratio calculations and survival curve plotting were performed using GraphPad Prism (version 10.3).

### Cell lines

All cell lines were obtained from the Cell Services facility of The Francis Crick Institute and verified as mycoplasma-free. All human cell lines were further validated by DNA fingerprinting.

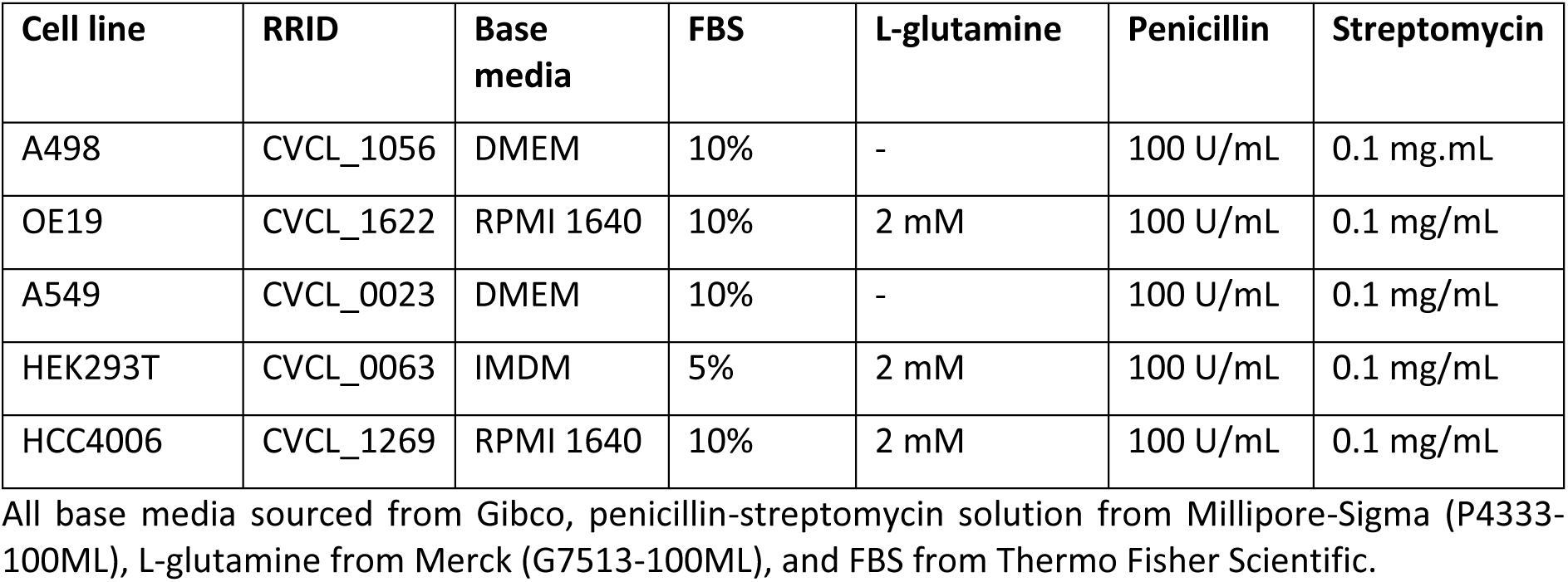

| Cell line | RRID | Base media | FBS | L-glutamine | Penicillin | Streptomycin |
| --- | --- | --- | --- | --- | --- | --- |
| A498 | CVCL_1056 | DMEM | 10% | - | 100 U/mL | 0.1 mg/mL |
| OE19 | CVCL_1622 | RPMI 1640 | 10% | 2 mM | 100 U/mL | 0.1 mg/mL |
| A549 | CVCL_0023 | DMEM | 10% | - | 100 U/mL | 0.1 mg/mL |
| HEK293T | CVCL_0063 | IMDM | 5% | 2 mM | 100 U/mL | 0.1 mg/mL |
| HCC4006 | CVCL_1269 | RPMI 1640 | 10% | 2 mM | 100 U/mL | 0.1 mg/mL |

All base media sourced from Gibco, penicillin-streptomycin solution from Millipore-Sigma (P4333-100ML), L-glutamine from Merck (G7513-100ML), and FBS from Thermo Fisher Scientific.

### Cell transfections

HEK293T and A549 cells were seeded at a density of 200,000 cells/well in 2ml of culture media 24 hours prior transfection in 6-well plates. Cells were then transfected with 5 µg of plasmid each expressing the following sequence (see table): CHRNA5 (pcDNA3.1-CHRNA5, Genewiz) or CHRNA5[AluSz] (pcDNA3.1-CHRNA5[AluSz], Genewiz) or CHRNA5-Full intron 5 (pcDNA3.1-CHRNA5-Full intron 5, Genewiz) or CHRNA5-RTE deleted (pcDNA3.1-CHRNA5-RTE deleted, Genewiz) using Lipofectamine 3000 transfection reagent (Thermo Fisher). Cell were then seeded for immunofluorescence staining.

### Reverse transcriptase-based quantitative PCR (RT-qPCR)

RNA was extracted using the RNeasy kit (Qiagen). cDNA was synthesized using the Maxima First Strand cDNA Synthesis Kit (Thermo Fisher), and qPCR performed using Applied Biosystems Fast SYBR Green (Thermo Fisher) using the following primers:

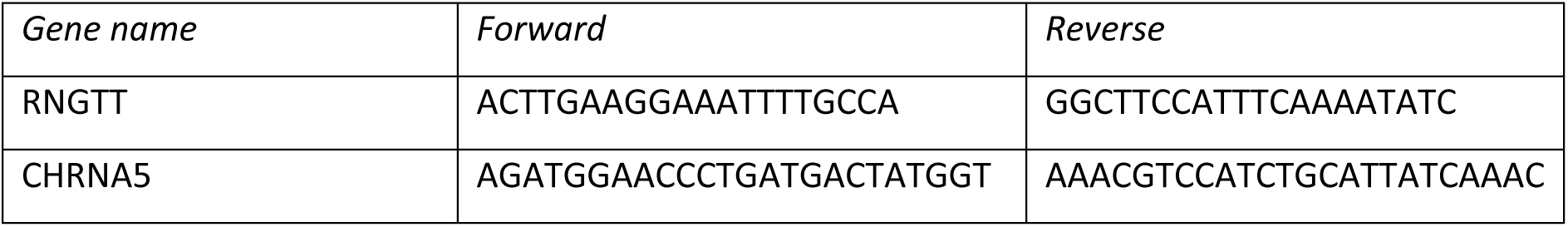
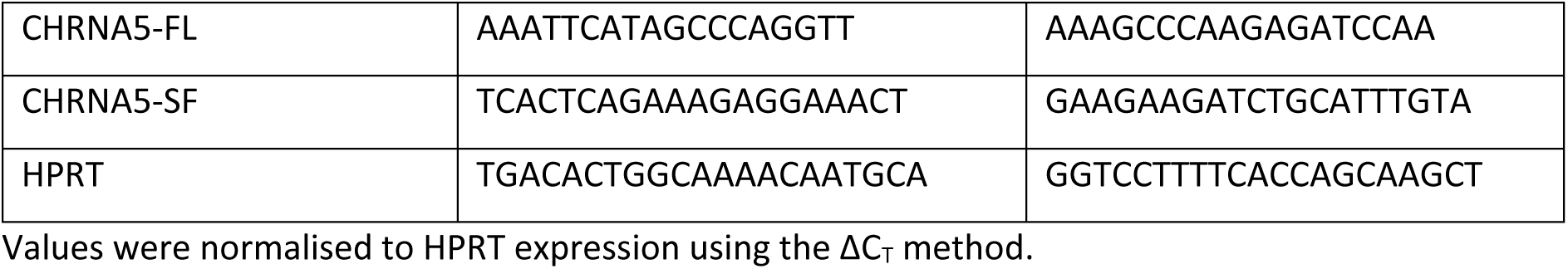

### PCR

PCR was performed using KOD Hot Start Master Mix (Sigma) with the following primers:

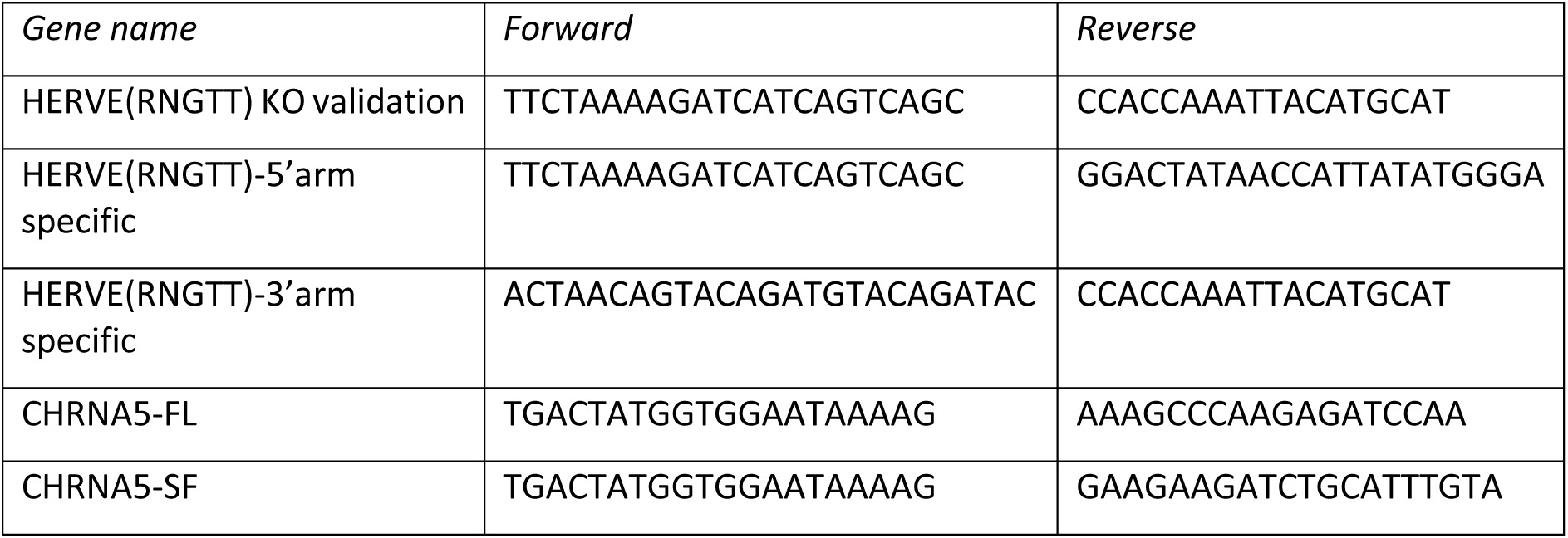

### Cas9-mediated editing

The HERVE provirus in the RNGTT locus was targeted by the following guide RNA (gRNA) sequences:

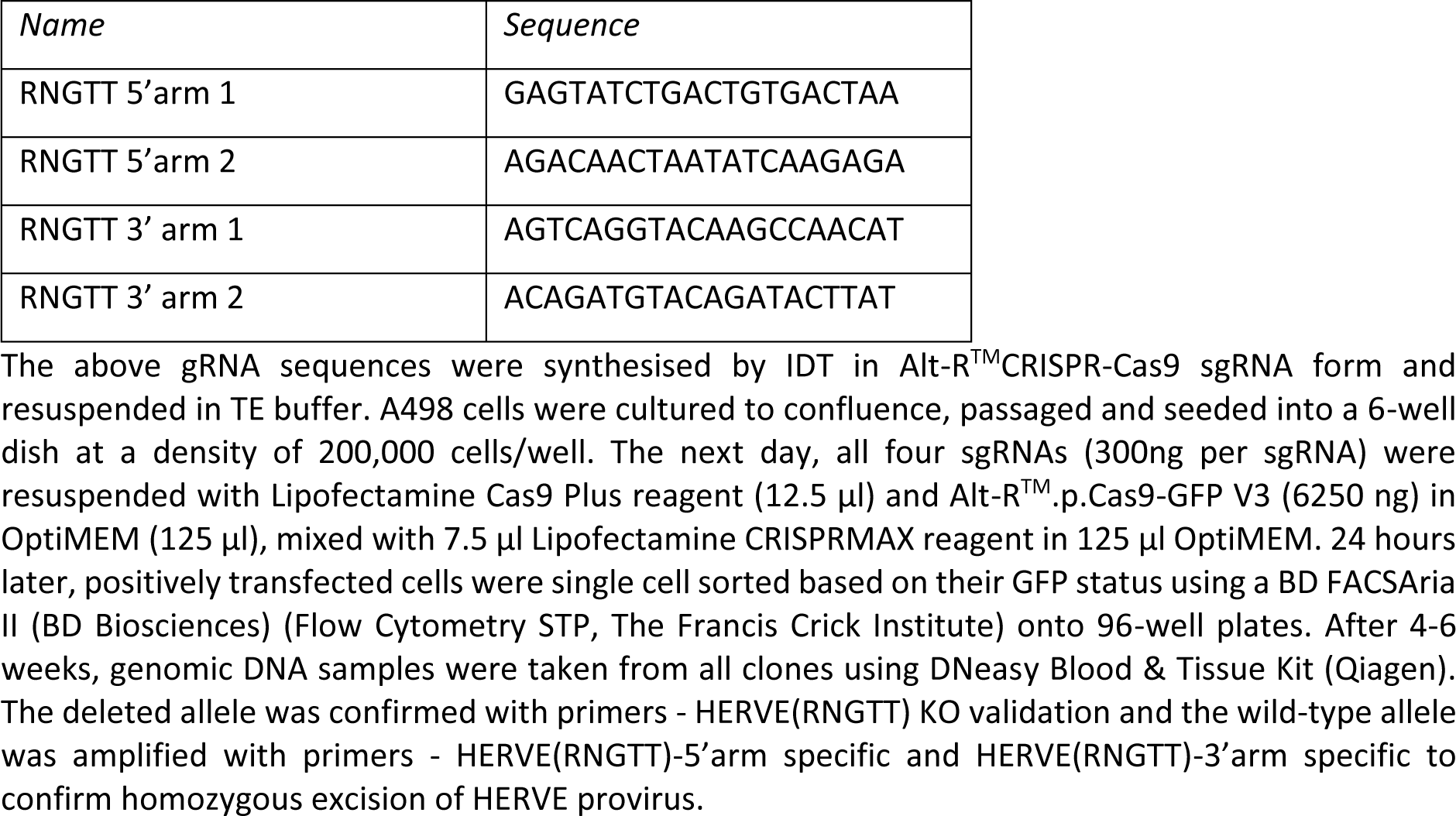

### Amplicon Sanger sequencing

Genomic DNA PCR products from A498 cells were sent to Genewiz for PCR clean-up and Sanger sequencing with the following sequencing primers:

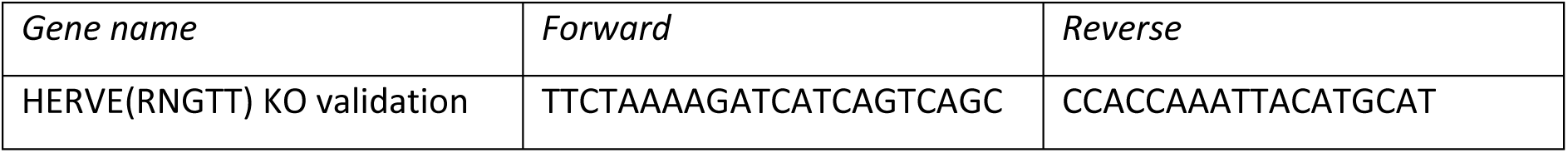

### Amplicon next-generation sequencing

Amplicons from HCC4006 cell cDNA specific for the *HERVH Xp22.2-AS* isoforms were amplified using the following primers:

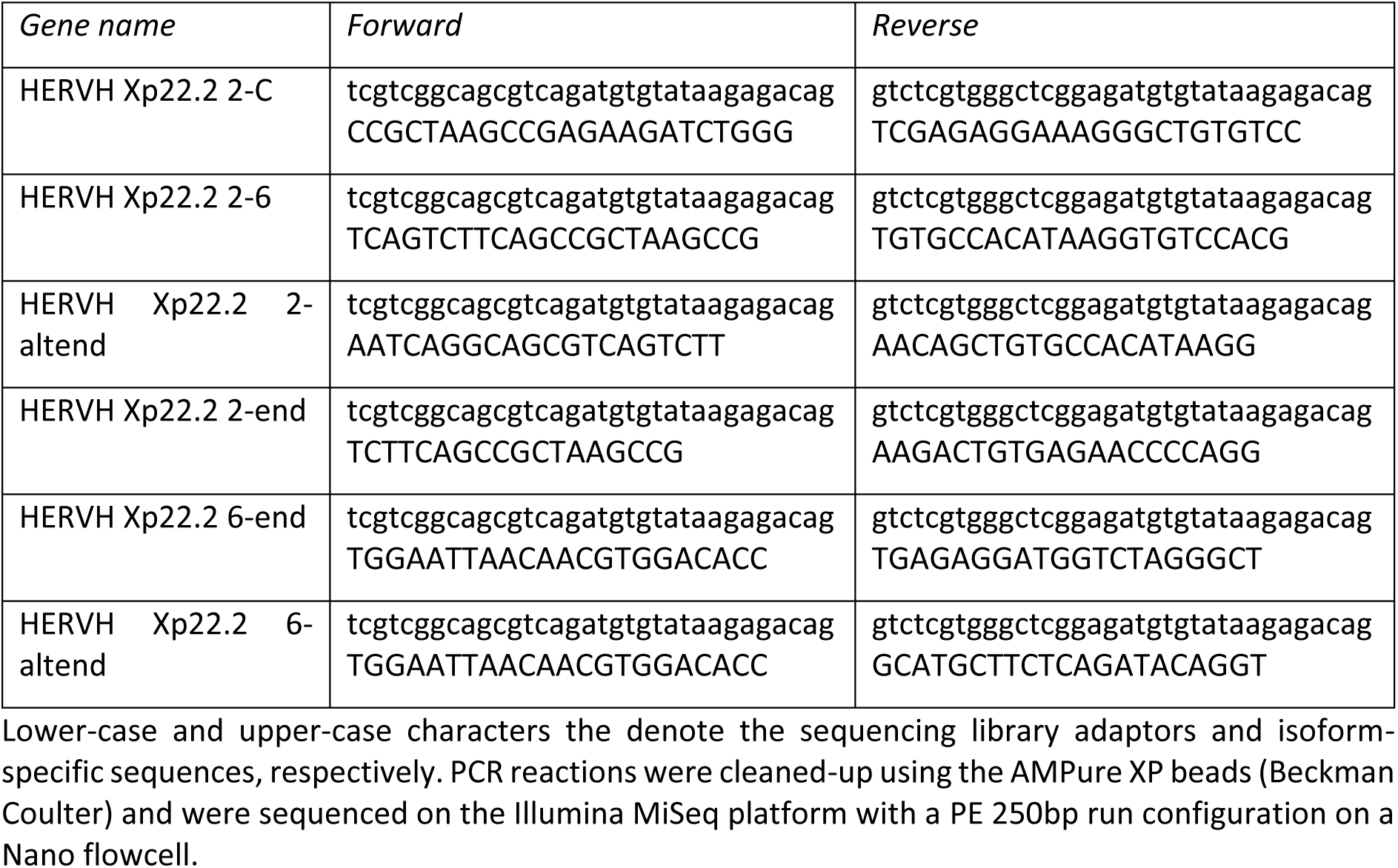

### Immunofluorescence

HEK293T cells transfected with pcDNA3.1-CHRNA5 or pcDNA3.1-CHRNA5[AluSz] were grown on 1.5mm coverglass dishes (MatTek, Cat #P35G-1.5-20-C) were fixed using 10% neutrally buffered formalin (Genta Medical) for 15 min. Non-specific staining of non-permeabilized cells was blocked with 1% bovine serum albumin (Sigma-Aldrich). Primary antibody incubation for HA-tag antibody (Santa Cruz Biotechnology, sc-7392) was performed overnight at 4°C at a 1:100 dilution. Secondary antibody incubation using Goat Anti-Mouse IgG H&L AlexaFluor594 Antibody (Abcam, Cat #ab150116) was carried out the next day for 1 hour at room temperature at a 1:1,000 dilution. Nuclear staining was performed using Hoechst 33342 (Thermo Fisher). Samples were imaged on the Zeiss Observer.Z1 (Carl Zeiss Meditec AG) using Micro-Manager 2.0 software.

### Retroviral transduction

Stably transduced cell lines were produced through viral infection and single cell sorting on green fluorescent protein (GFP) and mCherry using the MoFLO XDP cell sorter (BD Biosciences, Flow Cytometry Team, The Francis Crick Institute). Virus was generated using HEK293T cells plated at a density of 1.5×10^6^ cells per 60 mm well incubated with a mixture of serum-free IMDM (Sigma-Aldrich, I3390), GeneJuice (VWR International Ltd., 70967-4), and 5 μL of plasmid DNA. Plasmids used were vesicular stomatitis virus glycoprotein (VSVg) plasmid (pcVG-wt), pHIT60, and the open reading frames of the sequences of interest cloned into the pRV-IRES-GFP or pRV-IRES-mCherry vector (see table). Cloning the open reading frames into the vector was carried out Genewiz LLC, and was followed by sequencing to verify the plasmid structure. Virus-containing supernatant was collected and added to HEK293T cells (plated at a density of 8.5×10^4^ cells per 35 mm well) along with polybrene (Sigma-Aldrich, TR-1003-G), and spun at 1200 RPM for 45 minutes. After three days, populations were single cell sorted on GFP, or mCherry expression using a BD FACSAria II (BD Biosciences) (Flow Cytometry STP, The Francis Crick Institute).

### Protein preparation for Western Blot

Cells were washed twice with phosphate-buffered saline (PBS) stored at 4°C before being incubated on ice with radioimmunoprecipitation assay (RIPA, Sigma-Aldrich, R0278-50ML) buffer for 30 minutes to lyse the cells. The mixture was then spun at 14000 RPM for 10 minutes at 4°C. The protein concentration of the lysate was measured using the Pierce^TM^ BCA protein assay kit (Thermo Scientific, 23225). Stock solutions at a protein concentration of 500 μg/mL were made by mixing 100 μL of sample buffer (Laemmli 2x concentrate, Sigma-Aldrich, S3401-10VL), with 100 μg of protein lysate and RIPA buffer to a final volume of 200 μL. Stock solutions were heat denatured at 95°C for 5 minutes before being frozen at −20°C.

### Western Blot

Sample stock solutions were thawed on ice before being boiled at 95°C for 5 minutes. 10 μg of protein per sample was loaded into a 4–20% Mini-PROTEAN® TGX™ precast polyacrylamide gel (Bio-Rad, 4561094) alongside a protein ladder (PageRuler Plus Prestained Protein Ladder, 10 kDa to 250 kDa, ThermoFisher, 26619). The gel electrophoresis was run in a Mini-PROTEAN® Tetra Vertical Electrophoresis Cell (Bio-Rad) filled with protein running buffer (Media Team, The Francis Crick Institute) at 180 V for 40 minutes. Samples were transferred to a 0.2 μm nitrocellulose membrane (Trans-Blot Turbo Mini 0.2 μm Nitrocellulose Transfer Pack, Bio-Rad, 1704158) using the Trans-Blot Turbo dry transfer system (Bio-Rad, 1704150) turbo setting for mini TGX gels before blocking with 5% skimmed milk (Marvel) in Tris-buffered saline with 0.5% Tween-20 (TBS-T, Media Team, The Francis Crick Institute) for 1 hour. Membranes were stained overnight at 4°C with the anti-FLAG antibody (Sigma-Aldrich, F1804) diluted at 1:1000 in 5% skimmed milk in TBS-T. Membranes were washed with TBS-T at room temperature before the horseradish peroxidase (HRP)-conjugated anti-mouse secondary antibody (Abcam, ab6728**Error! Reference source not found.**) was added, diluted at 1:1000 in 3% skimmed milk in TBS-T and incubated at room temperature for 1 hour. Membranes were then washed in TBS-T and visualised by enhanced chemiluminescence using Clarity™ Western ECL Substrate (Bio-Rad, 1705060) on a ChemiDoc XRS+ (Bio-Rad).

### Transwell migration assay

Cell migratory capacity was examined by transwell migration assay that uses chemotactic gradient to assess how cells migrate through a porous membrane. Cells were trypsinised and resuspended in serum-free media. They were then seeded into Millicell Hanging Cell Culture Insert (Millipore, PTEP24H48) at a density of 20×10^3^ for A498 and A549 clones, while the lower chamber contained fresh culture media with 30% FBS. The cells were allowed to migrate for at least 48 hours. The inserts were washed with PBS and fixed with 10% neutrally buffered formalin (Genta Medical) for 10 minutes. Those cells that did not migrate were removed. The cells on the lower surface of the insert were washed with PBS again, stained with crystal violet solution (Sigma Aldrich, V5265). The cells on each insert were counted in 4 random fields under Zeiss Observer.Z1 (Carl Zeiss Meditec AG) using Micro-Manager 2.0 software.

### Cell growth assays

Growth and proliferation of A498 parental control (1E7), A498 *HERVE 6q15^-/-^* (2D11) clone, A549 parental and canonical *CHRNA5* and *CHRNA5[AluSz]* expressing A549 clones was assessed by real-time quantitative live-cell imaging using the Incucyte Live-Cell Analysis System (Sartorius). Cells were seeded into 96-well plates 24 hours prior measurement in Incucyte system at a density of 2000 and 1000 cells per well for A498 and A549, respectively. Image and confluency measurement were taken every 3 hours for at least 72 hours. Cell growth data for RNGTT-deficient cell lines in CCLE were downloaded from Dependency Map (DepMap) portal (https://depmap.org/portal) [23].

### Statistical analyses

Statistical comparisons were made using GraphPad Prism 10.3 (GraphPad Software), SigmaPlot 14.0, Qlucore Omics Explorer v3.9, or R (versions 3.6.1-4.0.0). Parametric comparisons of normally distributed values that satisfied the variance criteria were made by unpaired or paired Student’s *t*-tests or One Way Analysis of variance (ANOVA) tests with Bonferroni correction for multiple comparisons. Data that did not pass the variance test were compared with non-parametric two-tailed Mann-Whitney Rank Sum tests (for unpaired comparisons) or Kruskal-Wallis test with Dunn’s multiple comparisons correction.

## Results

### Widespread potential for RTE-mediate disruption of adjacent gene function

To identify cases where aberrant transcriptional inclusion of RTEs may affect local gene function, we searched for gene loci that exhibit a specific switch in RNA isoform expression. To this end, we used a cancer transcriptome assembly that captures diverse RTE-overlapping transcripts [8], and selected transcripts that were either upregulated or downregulated in a given cancer type compared with its respective normal tissue (Figure 1A-C). Intersection of the two lists identified transcripts that, despite exhibiting contrasting transcriptional behaviour (referred to here as discordant transcripts), were transcribed from the same locus (Figure 1A, B). For example, from a total of 225,544 transcripts differentially expressed between kidney renal clear cell carcinoma (KIRC) and normal kidney tissue, 62.9% were from loci that produced transcriptionally discordant RNA isoforms (Figure 1A-C), suggesting extensive changes in RNA isoform balance in this cancer type. Similar results were also obtained from analysis of colon adenocarcinoma (COAD), lung adenocarcinoma (LUAD), lung squamous cell carcinoma (LUSC) and esophageal adenocarcinoma (EAC), although the proportion of discordant transcripts was lower (11.3%-34.3%) in these cancer types (Figure 1C). Moreover, nearly half of the loci producing transcriptionally discordant RNA isoforms in KIRC were also found in at least one other cancer type, and this fraction was much higher (82.5%-92.2%) for the remaining types (Figure 1C). Compared with the entire previously assembled transcriptome, *Alu*, *MIR* and *L1* elements were particularly enriched in discordant transcripts from all types of cancer analysed, whereas LTR elements displayed a cancer type-specific pattern of enrichment (Figure 1D). These findings suggested extensive imbalances in RNA isoform expression in cancer through increased RTE utilisation and indicated common underlying mechanisms operating in multiple cancer types. Given the potential of alternative isoforms generated by RTE co-option to affect gene function or create new function, we next investigated specific effects on nearby or encompassing genes in detail.

**Figure 1.**
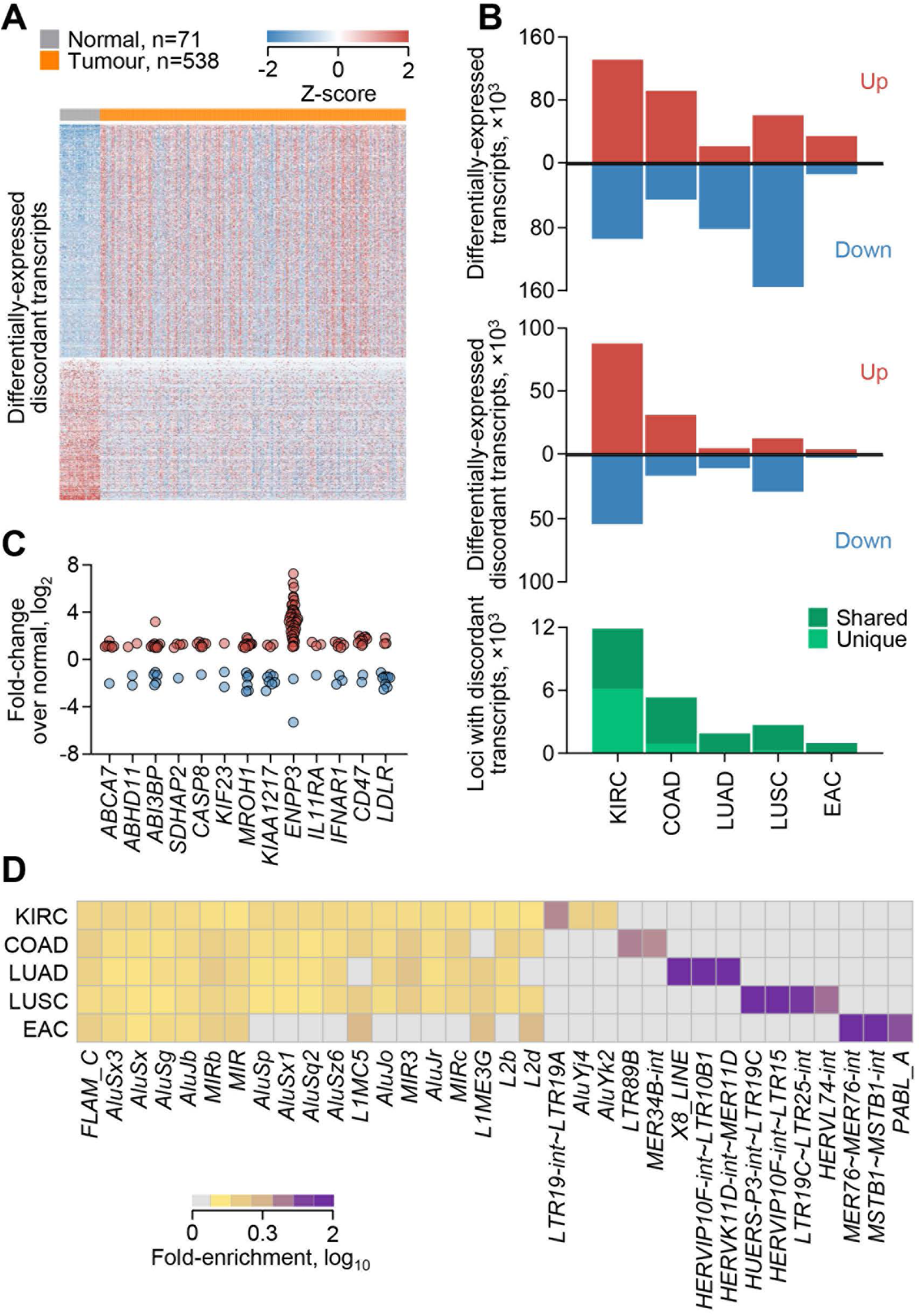
Widespread shifts in the balance of RNA isoform expression in cancer. **(A)** Heatmap of expression of 87,861 and 54,202 discordant transcripts that are upregulated and downregulated (≥1.5 fold-change, p≤0.05, q≤0.05), respectively, in TCGA KIRC samples, compared with normal kidney tissue, and overlap with loci producing transcripts in both categories. **(B)** Total number of differentially-expressed transcripts (≥1.5 fold-change, p≤0.05, q≤0.05) (*top*), differentially-expressed discordant transcripts (*middle*), and loci producing discordant transcripts (*bottom*) between the indicated cancer types and their respective normal tissue (KIRC n=538, normal kidney n=71; COAD n=239, normal colon n=39; LUAD n=419, normal lung n=24; LUSC n=362, normal lung n=24; EAC n=78, normal esophagus n=9). **(C)** Fold-change of individual discordant transcripts (symbols) overlapping with the indicated loci in KIRC samples. **(D)** Fold-enrichment for the indicated RTE subgroups in discordant transcripts, compared with the entire transcriptome. All indicated RTE subgroups were significantly enriched (p≤0.05).

### Downregulation of *RNGTT* transcription by an intronic *HERVE* integration

An example of a locus producing discordant transcripts in KIRC was *RNGTT*, encoding the mRNA capping enzyme with RNA 5’-triphosphate monophosphatase and guanylyltransferase activities. In its penultimate intron, *RNGTT* harbours a *HERVE* provirus, integrated in reverse orientation relative to *RNGTT* (Figure 2A). This *HERVE* provirus, referred to here as *HERVE 6q15*, is known to be highly expressed in KIRC and to encode immunogenic retroviral proteins, which contribute to tumour immune control [24–27]. However, the consequences of its transcriptional induction on *RNGTT* have not been previously considered. We found that transcription of *RNGTT* and the intronic *HERVE* provirus exhibited an inverse pattern in KIRC (Figure 2B-D). *HERVE 6q15* was not expressed in normal kidney tissue but was highly induced in ∼50% of KIRC cases, with significantly higher expression in later stages of the disease (Figure 2B, C). In contrast, *RNGTT* expression was significantly reduced in KIRC, compared with normal kidney tissue, and this reduction was more pronounced in cases with higher *HERVE 6q15* expression (Figure 2B, D), suggesting that transcriptional activation of the *HERVE* integration negatively impacted *RNGTT* expression. A similar negative correlation between *RNGTT* and *HERVE 6q15* transcription was also observed in RNA-seq data from individual renal cell carcinoma cell lines obtained from the Cancer Cell Line Encyclopedia (CCLE) (Figure 1E, F), where possible confounding effects of non-tumour cells in tumour samples can be excluded.

**Figure 2.**
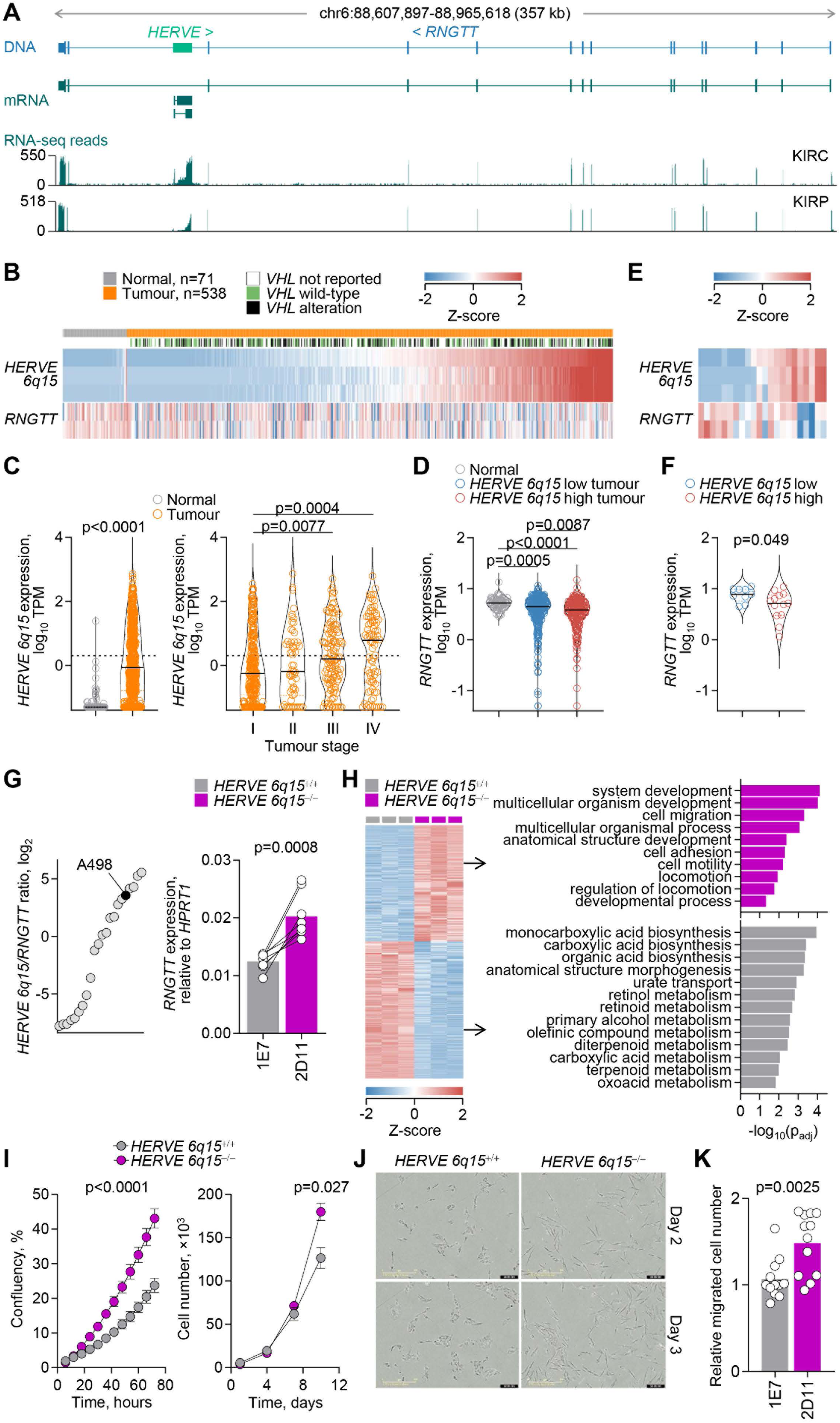
Effect of *HERVE* activation on *RNGTT* transcription. **(A)** Gene structure and integrated *HERVE* provirus, GENCODE-annotated and assembled transcripts, and RNA-seq traces of 24 combined KIRC and kidney renal papillary cell carcinoma (KIRP) samples at the *RNGTT* locus. **(B)** Expression of transcripts overlapping *HERVE 6q15* or the canonical *RNGTT* in normal kidney tissue and KIRC samples, ordered according to *HERVE 6q15* expression. **(C)** *HERVE 6q15* expression in TPM in the same samples as in b (p value calculated with Mann-Whitney test), and according to tumour stage (I n=271, II n=59, III n=123, IV n=82; p values calculated with Kruskal-Wallis test with Dunn’s multiple comparisons correction). **(D)** *RNGTT* expression (TPM) in normal kidney tissue (n=71) and in KIRC samples with low (n=322) and high (n=216) *HERVE 6q15* expression (p values calculated with Kruskal-Wallis test with Dunn’s multiple comparisons correction). **(E)** Expression of transcripts overlapping *HERVE 6q15* or the canonical *RNGTT* in renal cell carcinoma cell lines in CCLE, in columns ordered according to *HERVE 6q15* expression. **(F)** *RNGTT* expression (TPM) in renal cell carcinoma cell lines with low (n=9) and high (n=13) *HERVE 6q15* expression (p value calculated with two-tailed Student’s t-test). **(G)** Ratio of *HERVE 6q15* to *RNGTT* expression in renal cell carcinoma cell lines in CCLE (*left*), and *RNGTT* expression (assessed by RT-PCR and plotted relatively to *HPRT1* expression) in A498 cells with (clone 1E7, *HERVE 6q15*^+/+^) or without (clone 2D11, *HERVE 6q15*^-/-^) the *HERVE 6q15* provirus (*right*). Symbols represent replicates of 4 independent experiments, connected with lines (p value calculated with two-tailed paired Student’s t-test). **(H)** Heatmap of differential gene expression (≥2 fold-change, p≤0.05, q≤0.05) of between *HERVE 6q15*^+/+^ and *HERVE 6q15*^-/-^ A498 cells (*left*). Columns represent independent replicates. Gene ontology (GO) functional annotation of the differentially expressed genes (*right*) (p values calculated with g:Profiler using hypergeometric distribution tests and adjusted for multiple hypothesis testing using the g:SCS (set counts and sizes) algorithm, integral to the g:Profiler server (https://biit.cs.ut.ee/gprofiler)). **i**, Mean confluency (±SD, n=6 from 1 experiment) (*left*) and mean cell number (±SEM, n=3 from 1 experiment) (*right*) of *HERVE 6q15*^+/+^ and *HERVE 6q15*^-/-^ A498 cell cultures over time (p values calculated by two-tailed Student’s t-test of the area under the curve (AUC) values and by two-tailed Student’s t-test for day 10, respectively). **(J)** Representative images of of *HERVE 6q15*^+/+^ and *HERVE 6q15*^-/-^ A498 cell morphology 2 or 3 days after plating. **(K)** *In vitro* migration of *HERVE 6q15*^+/+^ (1E7) and *HERVE 6q15*^-/-^ (2D11) A498 cells. Symbols represent independent measurements (n=12, 4 fields of view from 3 independent experiments; p value calculated with two-tailed Student’s t-test).

In agreement with prior reports [25], *HERVE 6q15* transcription in KIRC was directly correlated with the degree of hypoxia (Figure S1A, B), although the strength of this correlation was likely affected by the ability of current hypoxia scores to reflect the pseudohypoxic state of this cancer type [20]. However, a direct effect of hypoxia on *HERVE 6q15* transcription was evident by analysis of RNA-seq data from renal cell carcinoma RCC4 cells with restored expression of the von Hippel-Lindau (VHL) tumour suppressor [28] (GSE120887) (Figure S1C, D). Hypoxia significantly increased *HERVE 6q15* expression in VHL-expressing RCC4 cells, but did not affect levels of *RNGTT* transcription, which were already very low and substantially lower than those of *HERVE 6q15* in these cells (Figure S1C, D).

The enzymatic activities encoded by *RNGTT* play an indispensable role in RNA capping, by catalysing the first step of a complex series of reactions leading to the addition of the methyl-7-guanosine cap on the 5’ ends of nascent RNAs [29]. In turn, RNA capping affects all subsequent aspects of RNA processing and function, including translation potential [29]. Accordingly, *RNGTT* is a common essential gene, required for growth of virtually all cancer cell lines (Figure S2), and its direct pro-tumour activity is associated with worse prognosis on most cancer types [29–31].

Given that high levels of *HERVE 6q15* transcription in most renal cell carcinoma cell lines may have reduced *RNGTT* transcription to levels that cannot be further reduced without compromising cell viability (Figure S2), we investigated a direct effect of *HERVE 6q15* on *RNGTT*, with the reverse experiment. We selected renal cell carcinoma A498 cells, which express a high *HERVE 6q15* to *RNGTT* ratio (Figure 1G), and used CRISPR/Cas9 editing to remove the *HERVE* provirus, together with two immediately adjacent *L1* integrations (ED. Figure 3). Assessed by RT-qPCR and compared with the *HERVE 6q15^+/+^* clone (1E7), expression of *RNGTT* was upregulated by ∼63% in the *HERVE 6q15^-/-^* clone (2D11) (Fig, 1G). *RNGTT* upregulation following *HERVE 6q15* deletion in A498 cells was accompanied by extensive transcriptional changes, with upregulation of genes involved in cell adhesion and migration, and downregulation of genes involved in metabolic processes (Figure 1H). Consistent with their transcriptional profile and the pro-tumour activities of *RNGTT*, *HERVE 6q15^-/-^* 2D11 cells exhibited increased *in vitro* growth, altered morphology and increased migration (Figure 1I-K). Lastly, supporting opposing transcriptional profiles, *RNGTT* and *HERVE 6q15* levels also showed the opposite correlation with KIRC survival, which was however an indirect result of *HERVE 6q15* association with later stages of the disease (Figure S4). Together, these results support a model of *HERVE 6q15* transcriptional induction during KIRC progression, which reduces, but does not abolish pro-tumour *RNGTT* expression.

**Figure 3.**
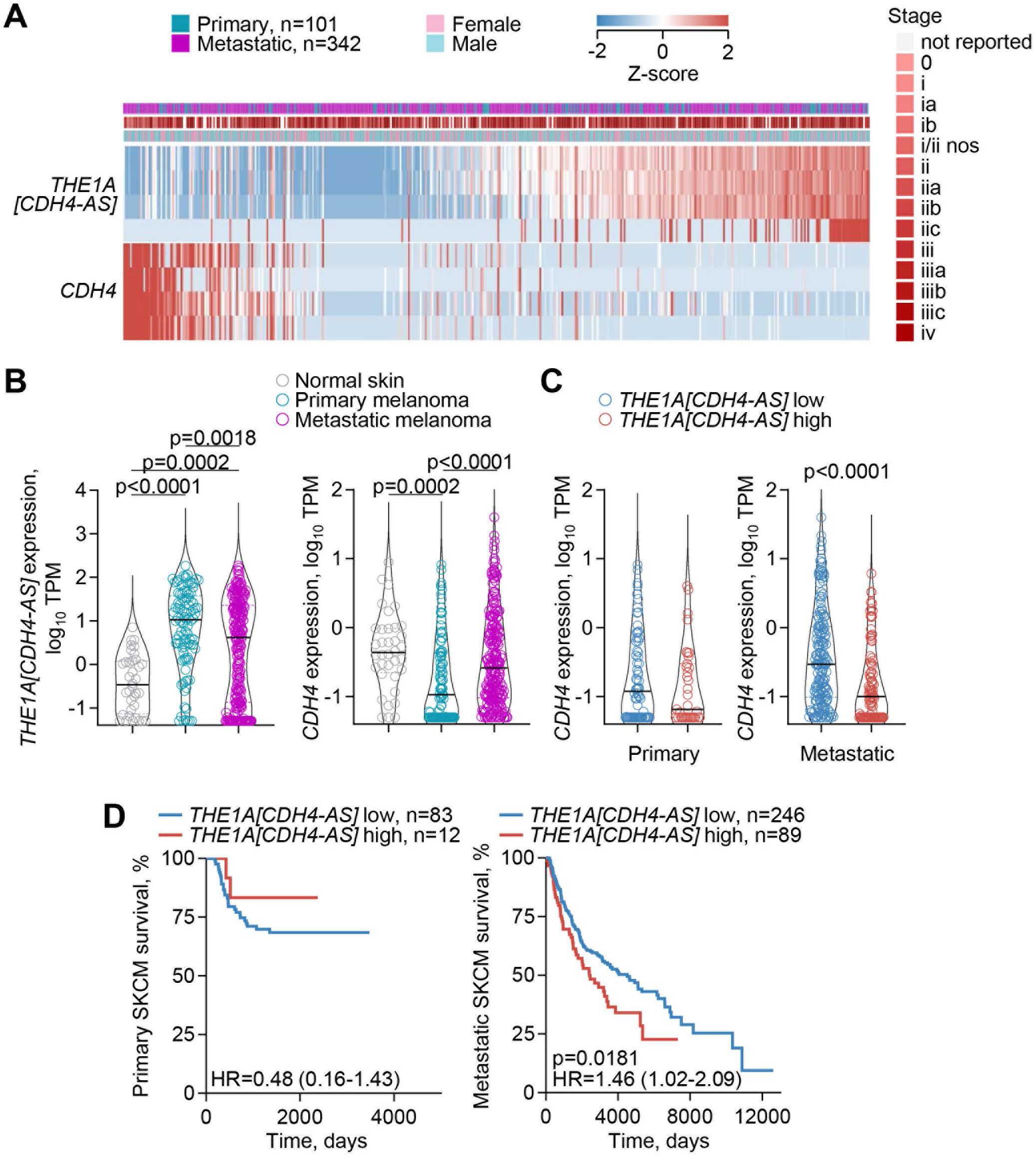
Effect of *THE1A* activation on *CDH4* transcription. **(A)** Expression of transcripts overlapping *THE1A[CDH4-AS]* or the canonical *CDH4* in primary and metastatic SKCM samples. **(B)** *THE1A[CDH4-AS]* and canonical *CDH4* expression (TPM) in normal skin tissue (n=36) and in primary (n=101) and metastatic (n=224) SKCM samples (p values calculated with Kruskal-Wallis tests with Dunn’s multiple comparisons correction). **(C)** *CDH4* expression (TPM) in primary and metastatic SKCM samples with low (n=66 and n=161, respectively) and high (n=35 and n=89, respectively) *THE1A[CDH4-AS]* expression (p value calculated with Mann-Whitney test). **(D)** Overall survival of primary and metastatic SKCM patients, stratified by *THE1A[CDH4-AS]* expression (p values calculated with log-rank tests).

### Regulation of Cadherin 4 by *THE1A*-driven antisense transcription

We have previously identified an antisense transcript, initiated by a *THE1A* retroelement and referred to as *THE1A[CDH4-AS]*, which is spanning two introns of the *CDH4* gene and expressed specifically in cutaneous and uveal melanomas [8]. To examine a possible effect of transcriptional activation of the intronic *THE1A* element on *CDH4* transcription we correlated the two in RNA-seq data from The Cancer Genome Atlas (TCGA) skin cutaneous melanoma (SKCM) samples (Figure 3). This analysis revealed a pattern of mutual exclusivity between *THE1A[CDH4-AS]* and *CDH4* transcription in both primary and metastatic SKCM, irrespective of disease stage (Figure 3A). However, compared with normal skin samples, the transcriptional activation of the *THE1A* element and concomitant downregulation of *CDH4* transcription were considerably more pronounced in primary than in metastatic disease (Figure 3B, C). Indeed, whereas significantly reduced in primary melanoma, overall *CDH4* transcription remained high in metastatic melanoma, particularly in samples with low *THE1A[CDH4-AS]* transcription (Figure 3B, C).

*CDH4* encodes Cadherin 4, also known as R-cadherin (retinal), a member of the calcium-dependent adhesion molecule superfamily, important in forming adherens junctions and in organ development [32, 33]. Cadherin-regulated cellular adhesion and detachment also determines migration and invasiveness of tumour cells and, consequently, the ability of several cancer types to metastasise [34, 35]. In the skin, E-cadherin (epidermal) and P-cadherin (placental) mediate heterotypic adhesion of neural crest-derived melanocytes with the surrounding epithelial cells [36]. Loss of E-cadherin and P-cadherin, and gain of N-cadherin (neuronal), known as a cadherin switching [35], facilitates melanoma cell metastasis and is associated with worse prognosis of SKCM [37]. While, its role in melanoma is less well studied, R-cadherin mediates adhesion with N-cadherin [32] and its overexpression in epidermoid carcinoma A431 cells causes the loss of E-cadherin and P-cadherin, through competition for the intracellular adaptor proteins catenins [38]. An involvement of R-cadherin in cadherin switching is further supported by reports of an essential role in tumorigenesis and metastasis in human osteosarcoma [39] and in a murine model of glioma [40], and of a tumour suppressive function in human colorectal and gastric cancers [41].

Consistent with a role for *CDH4* in the metastatic process suggested by findings in other cancer types, we found that transcriptional co-option of *THE1A[CDH4-AS]* specifically in SCKM, differentiates primary and metastatic disease and is differentially associated with survival in the two subtypes (Figure 3D). This association indicated that *THE1A[CDH4-AS]*-mediated suppression of *CDH4* in primary melanomas restrains their metastatic potential, whereas the failure to establish or the loss of such transcriptional effect facilitates metastasis.

### *HERVH*-driven downregulation of endosomal single-stranded RNA sensors TLR7 and TLR8

The *TLR7* and *TLR8* paralogue genes are arranged in tandem on chromosome Xp22.2, followed by a *HERVH* provirus, referred to here as *HERVH Xp22.2* (Figure 4A). Our assembly identified a number of antisense transcripts, initiated by the bidirectional promoter activity of the *HERVH Xp22.2* provirus, using a total of 10 alternative exons spread throughout the locus and extending to the upstream gene *PRPS2* (Figure 4A). Some of the assembled transcripts partially matched the annotated *TLR8-AS1* transcripts in GENCODE 46 (ENST00000451564) and RefSeq (NR_030727), which appeared incomplete (Figure 4A). Splice junction analysis of RNA-seq data from TCGA LUAD samples revealed two main groups of *HERVH Xp22.2*-initated antisense transcripts (collectively referred to as *HERVH Xp22.2-AS*), one terminating between the *TLR7* and *TLR8* loci and one terminating within the *PRPS2* locus (Figure 4A). To validate the complex splicing pattern, we amplified the corresponding cDNAs from key splicing isoforms expressed in lung adenocarcinoma HCC4006 cells. Deep-sequencing of the amplicons confirmed the alternative use of middle and terminal exons, as well as the balance of shorter and longer *HERVH Xp22.2-AS* isoforms (Figure 4A).

**Figure 4.**
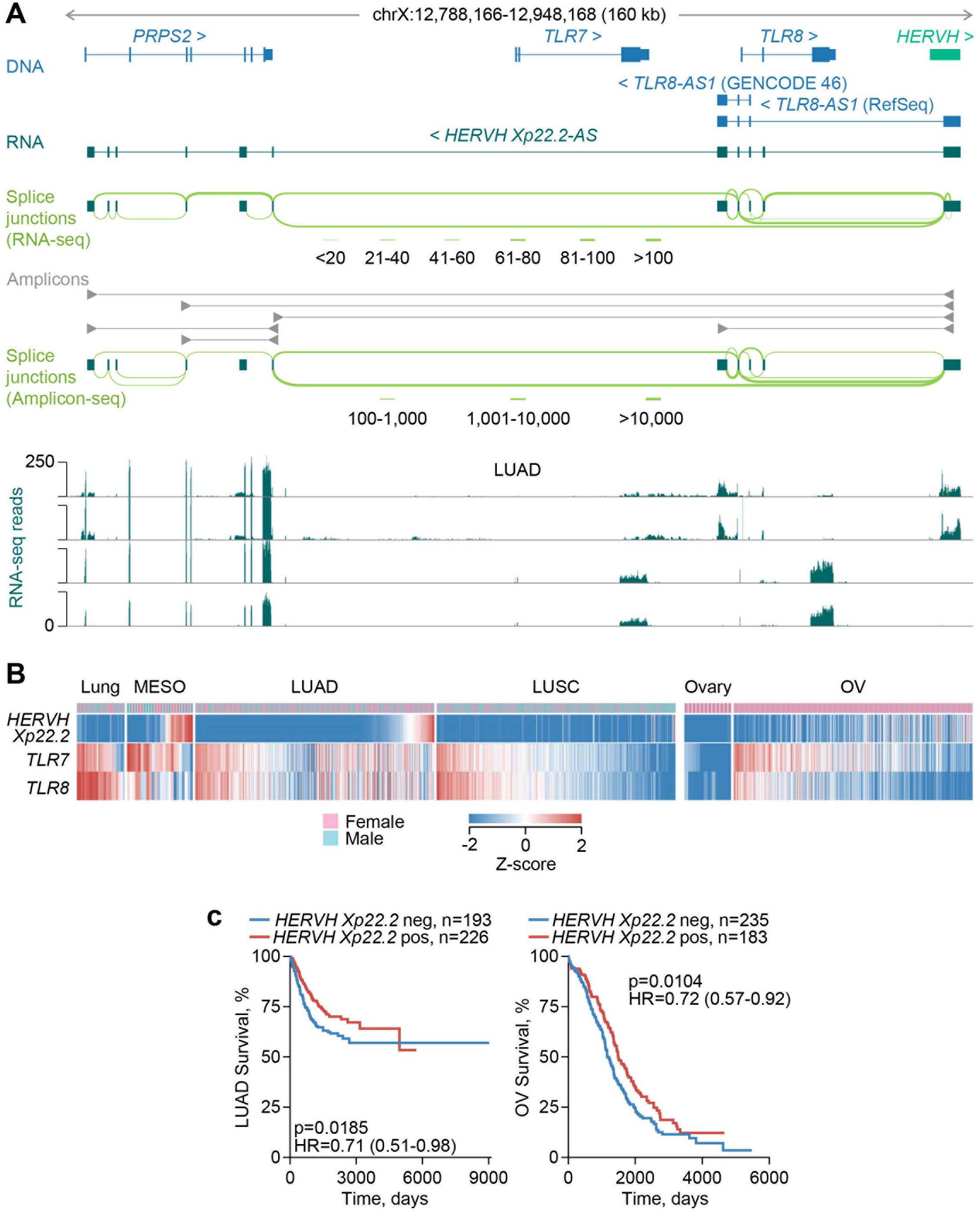
Effect of *HERVH* activation on *TLR7* and *TLR8* transcription. **(A)** Gene structure and integrated *HERVH* provirus at the *PRPS2-TLR7-TLR8* locus, GENCODE- and RefSeq-annotated and assembled transcripts, splice junction analysis of LUAD RNA-seq data, amplicons used for transcript validation, splice junction analysis of amplicon sequencing, and RNA-seq traces of LUAD samples with (*top two*) or without (*bottom two*) *HERVH* transcriptional activation. **(B)** Expression of transcripts overlapping *HERVH Xp22.2* or the canonical *TLR7* or *TLR8* in normal lung (n=36) and ovary (n=12) tissue, and in MESO (n=24), LUAD (n=419), LUSC (n=362) and OV (n=419) samples. **(C)** Overall survival of LUAD (*left*) and OV (*right*) patients, stratified by *HERVH Xp22.2* expression (p values calculated with log-rank tests).

Across several cancer types and respective normal tissues, *HERVH Xp22.2-AS* was transcriptionally activated in a substantial proportion of samples from ovarian serous cystadenocarcinoma (OV), mesothelioma (MESO) and testicular germ cell tumours (TGCT), and a smaller proportion of samples from other cancers, including LUAD and LUSC, but remained inactive in normal tissues, with the possible exception of a small number of EBV-transformed B cell lines (Figure S5A). Notably, *HERVH Xp22.2-AS* transcription was strongly anti-correlated with *TLR7* and *TLR8* transcription is cancers where it was expressed (Figure 4A, B; Extended Data Fig, 5B). Indeed, whereas normal lung tissue expressed *TLR7* and *TLR8* highly and proportionally, without detectable *HERVH Xp22.2-AS* expression, ∼13% of LUAD samples expressed high levels of *HERVH Xp22.2-AS*, but not of *TLR7* and *TLR8* (Figure S5B), indicating an inhibitory effect of *HERVH Xp22.2-AS* transcriptional activation on sense transcription. This effect appeared to extend also to the *PRPS2* locus, which was expressed only in LUAD samples, but not in normal lung tissue, and exhibited a significant anti-correlation with *HERVH Xp22.2-AS* expression (Figure S5B). Similar results were obtained also in LUSC, as well as MESO and OV, where *HERVH Xp22.2-AS* was highly expressed and *TLR7* and *TLR8* were downregulated in nearly half of the cases (Figure 4B).

TLR7 and TLR8 are endosomal sensors of single-stranded RNA [42] that play essential roles in the defence against viral infection and in the induction of B cell systemic autoimmunity [43]. Recent studies have indicated an essential, yet dual role for TLR7 and TLR8 also in cancer progression and immune control [44, 45]. Whereas their ligation in immune cells may enhance anti-tumour activity, TLR7 and TLR8 are also expressed in tumour cells where the mediate a pro-tumour effect [44, 45]. Indeed, tumour cell-intrinsic expression of TLR7 and TLR8 promotes their growth and survival *in vitro* [46, 47] and is associated with poor clinical outcome in non-small cell lung carcinomas (NSCLC) [48–50]. This pro-tumour effect of TLR7 is further supported by studies in animal models [48].

A pro-tumour effect of tumour cell-intrinsic TLR7 and TLR8 expression would predict that their downregulation by *HERVH Xp22.2-AS* transcriptional activation has an anti-tumour effect. Although survival analyses of *TLR7* and *TLR8* expression are confounded by their expression both in tumour cells and in immune cells, the strict tumour specificity of *HERVH Xp22.2-AS* expression reflects tumour cell-intrinsic transcriptional states. Indeed, we found that higher *HERVH Xp22.2-AS* expression in tumour samples is significantly associated with better prognosis in both LUAD and OV (Figure 4B), consistent with a protective effect of *HERVH Xp22.2-AS* transcriptional activation.

### Downregulation of *APOBEC3B* expression by *MER11C* element co-option

Similar to *TLR7* and *TLR8* paralogues, members of the Apolipoprotein B editing complex 3 (APOBEC3) family of enzymes are encoded by genes arranged in a cluster on chromosome 22 (Figure 5A, B). They catalyse cytidine deamination in DNA or RNA substrates, which can potently inhibit virus and RTE replication, but can also drive genomic diversity and instability in cancer [51, 52]. Current evidence implicates APOBEC3A and APOBEC3B as the two enzymes primarily responsible for the mutational signatures in human cancers and indicate a role for APOBEC3B in the regulation of APOBEC3A [53]. Expression of human APOBEC3B in mice enhances their susceptibility to tumours and also causes male infertility [54], supporting a pro-tumour role, as well as a detrimental effect on the genetic integrity of the male germline.

**Figure 5.**
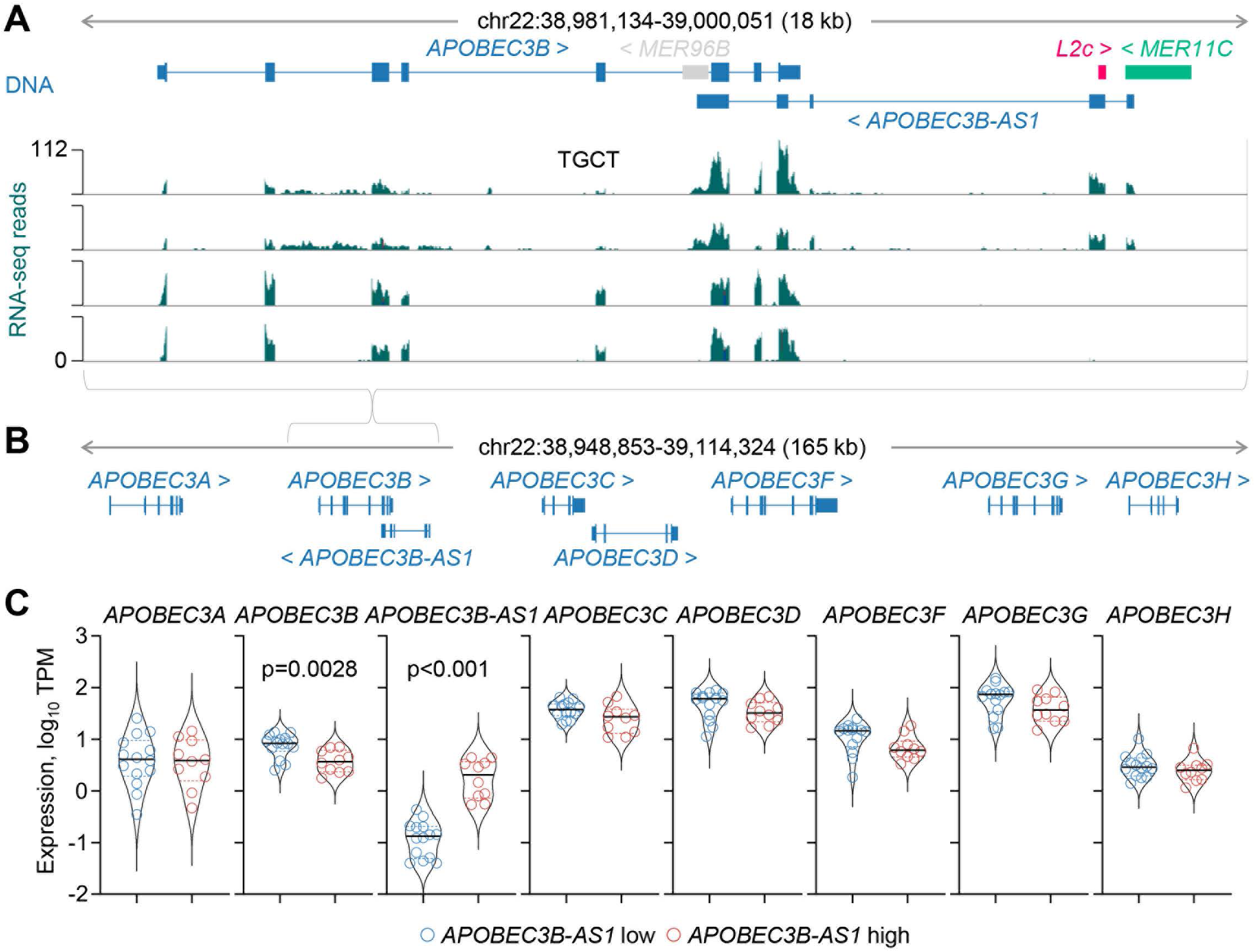
Effect of *MER11C* activation on *APOBEC3B* transcription. **(A)** Gene structure and integrated *MER11C, L2c* and *MER96B* RTEs at the *APOBEC3B* locus, and RNA-seq traces of TGCT samples with (*top two*) or without (*bottom two*) *MER11C* transcriptional activation. **(B)** Gene structure at the extended *APOBEC3* gene cluster. **(C)** Expression (TPM) of *APOBEC3B-AS1* and the indicated *APOBEC3* genes in TGCT samples with low (n=14) and high (n=10) *APOBEC3B-AS1* expression (p values calculated with Mann-Whitney test and Student’s t-test for *APOBEC3B-AS1* and *APOBEC3B*, respectively).

In this locus, we have identified a transcript matching annotated transcript *APOBEC3B-AS1* (ENST00000513758) and transcribed in the reverse orientation in relation to the *APOBEC3* genes (Figure 5A, B). This transcript was initiated by a *MER11C* element integrated between the *APOBEC3B* and *APOBEC3C* genes and extended over the first 3 *APOBEC3B* exons (Figure 5A, B). High *APOBEC3B-AS1* expression was highly specific to TGCT samples, with very low expression in other cancer types or normal tissues (Figure S6). Importantly, transcriptional activation of the *MER11C* element in TGCT samples was accompanied by reduction specifically in transcription of the overlapping *APOBEC3B* gene, whereas transcription of all other *APOBEC3* genes in this cluster remained unaffected (Figure 5C).

### Reduction of ENPP3 potential by a switch to non-functional isoforms

The *ENPP3* locus produced multiple discordant transcripts, the majority of which were highly upregulated (Figure 1B). In addition to *ENPP3*, the locus also contains two other annotated genes, *CTAGE9* and *OR2A4*, both located in intron 16 of *ENPP3* and transcribed in the reverse orientation (Figure 6A). Inspection of the locus identified several novel isoforms created by transcriptional inclusion of RTEs (Figure 6A). These included a transcript using *L2a* and *AluSx* elements (*ENPP3[L2a/AluSx]*) also in intron 16 as alternative terminal exon and polyadenylation site (Figure 6A). They also included a transcript using *AluSx3* and *L2a* elements as alternative second and terminal exons, respectively (*ENPP3[AluSx3/L2a]*), partially matching annotated GENCODE 46 transcript ENST00000427707 (Figure 6A). Two additional transcripts were created by the use of a *PABL_B* element as alternative promoter (*[PABL_B]ENPP3*) or an *AluSg* element as alternative terminal exon (*ENPP3[AluSg]*) (Figure 6A). Compared with normal kidney tissue, expression of *ENPP3* was found significantly elevated in KIRC samples, in agreement with prior reports [55, 56], as was expression of alternative *ENPP3* isoforms, as well as of *CTAGE9* and an *L1MDa* integration straddling *CTAGE9*, whereas expression of *OR2A4* was similar (Figure 6B; Figure S7A). Alternative *ENPP3* isoform expression accounted for a substantial proportion (∼32%) of total *ENPP3* transcription, with stable balance through progressive stages of the disease, and was primarily driven by the *ENPP3[L2a/AluSx]* and *ENPP3[AluSx3/L2a]* transcripts (Figure 6B; Figure S7B). Notably, whereas other *ENPP3* isoforms were expressed proportionally with the canonical isoform, expression of *ENPP3[L2a/AluSx]* and *CTAGE9* did not follow this pattern and appeared to be independently regulated (Figure 6C).

**Figure 6.**
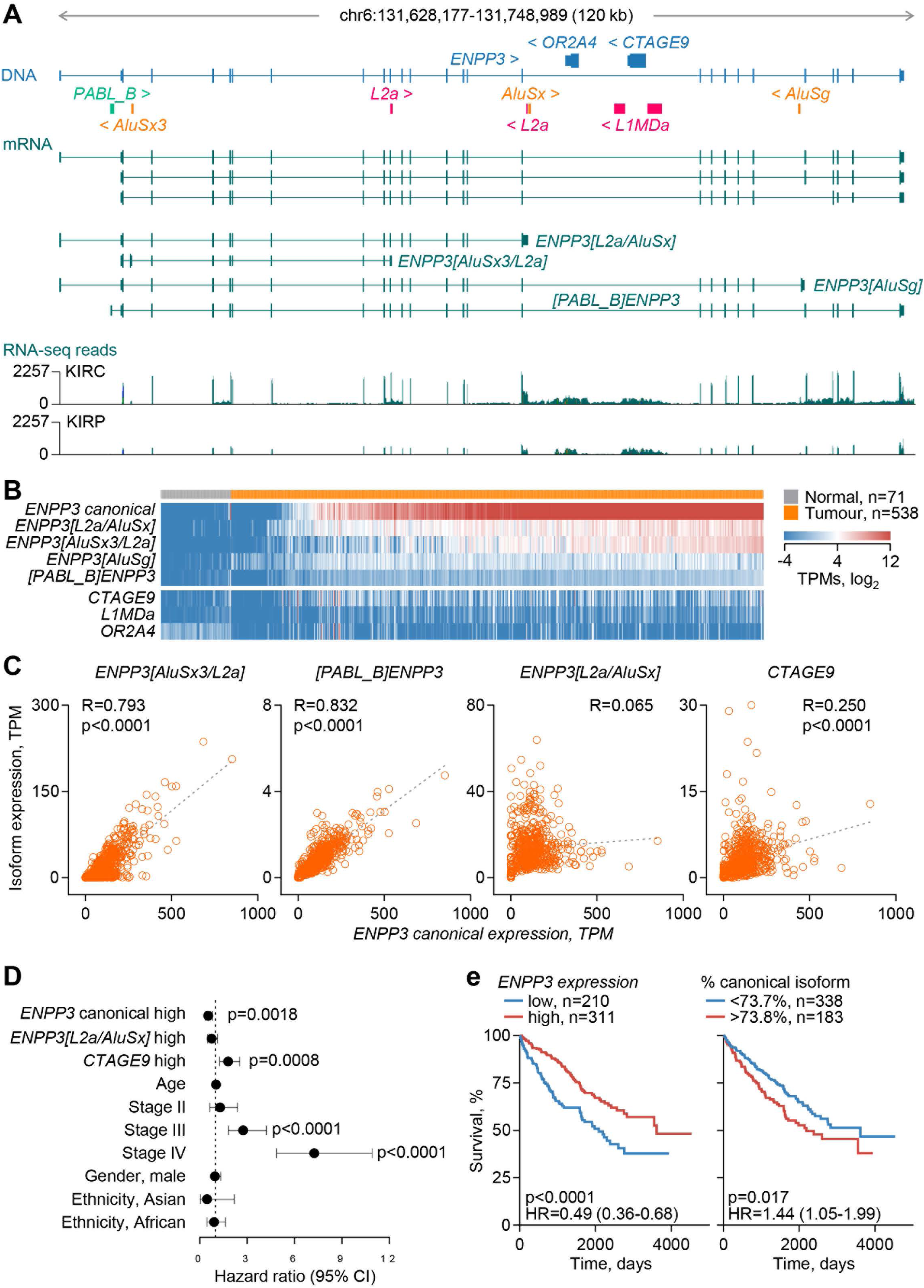
Effect of local RTEs on *ENPP3* functional and non-functional isoform balance. **(A)** Gene structure and exonised RTEs, GENCODE-annotated and assembled transcripts, and RNA-seq traces of 24 combined KIRC and KIRP samples at the *ENPP3* locus. **(B)** Expression of transcripts overlapping the indicated *ENNP3* isoforms or overlapping genes and RTEs in normal kidney tissue and KIRC samples, ordered according to canonical *ENPP3* expression. **(C)** Correlation of *ENPP3* isoform and *CTAGE9* expression (TPM) in KIRC samples (n=538) (p values calculated with linear regression). *CTAGE9* expression is capped at 30 TPM. **(D)** Overall survival hazard ratios (HRs) for the indicated variables in KIRC patients (*ENPP3* canonical high n=311, reference low n=210; *ENPP3[L2a/AluSx]* high n=214, reference low n=307; *CTAGE9* high n=174, reference low n=347; Age n=521; Stage II n=56, III n=123, IV n=82, reference I n=257; Gender, male n=338, reference female n=183; Ethnicity, Asian n=8, African n=55, reference White n=450). Error bars represent 95% CIs (p values calculated with Cox proportional hazards regression). **(E)** Overall survival of KIRC patients, stratified by *ENPP3* expression (*left*) or the fraction of the canonical *ENPP3* isoform in total *ENPP3* expression (*right*) (p values calculated with log-rank tests).

ENPP3, also known as CD203c, is a type II transmembrane protein that catalyses the hydrolysis of extracellular nucleotides [57]. It was originally identified as a basophil and mast cell activation marker, regulating allergic inflammation by hydrolysing extracellular ATP [57, 58]. More recently, ENPP3 has also been implicated in the regulation of extracellular levels of cGAMP (2′3′-cyclic guanosine monophosphate), a second messenger for the activation of the STING (stimulator of interferon genes) pathway and the production of type I IFNs in viral infection and cancer [59]. Supporting a pro-tumour role, loss of ENPP3 cGAMP hydrolase activity in mice renders them more resistant to primary tumour growth and metastasis [59]. In addition to regulating the tumour immune environment, cell-intrinsic expression of *ENPP3* has been reported to promote cell migration [60] and to be essential for the growth of renal cell carcinoma cell lines [56], further supporting a pro-tumour function. We, therefore, considered the potential activity of the alternative ENPP3 isoforms.

Exonisation of the *AluSx3* element after the first coding exon of in the *ENPP3[AluSx3/L2a]* isoform creates a premature termination after codon 53, producing a severely truncated product (UniProt ID: E7ETI7), unlikely to retain any function. The *ENPP3[L2a/AluSx]* isoform has the potential to produce a larger protein, retaining the transmembrane helix and most of the phosphodiesterase domain, but missing the nuclease domain (Extended Data F. 8A), which could exert altered enzymatic activities. However, in contrast to the canonical isoform, which was readily detectable upon overexpression in HEK293T cells, the *ENPP3[L2a/AluSx]* isoform failed to produce a product of the expected or higher mass, indicative of protein instability (Figure S8B). These results suggested that alternative *ENPP3* isoforms are non-functional and their production is, therefore, at the expense of the canonical, thereby compromising the maximum capacity of the locus to produce the ENPP3 enzymatic activity. In turn, the reduction in ENPP3 potential could impede tumour progression. In multivariate analyses, overall *ENPP3* transcription was associated with favourable outcome in KIRC, as previously reported [55, 56], whereas *CTAGE9*, which encodes the cutaneous T cell lymphoma-associated antigen 9, showed the inverse association (Figure 6D). Pertinently, a low proportion of canonical *ENPP3* among *ENPP3* isoforms was significantly associated with better survival in KIRC (Figure 6E), supporting a model where the degree of switching to non-functional *ENPP3* isoforms through RTE co-option correlated with disease outcome.

### Loss of tumour cell-intrinsic *CHRNA5* function by *Alu* exonisation

*CHRNA5* encodes the alpha 5 subunit of heteropentameric nicotinic acetylcholine receptor (nAChR) complexes, which initiate signalling cascades upon ligand-gated ion influx [61]. Multiple studies have linked a genetic variant in *CHRNA5* exon 5 (rs16969968) with susceptibility to lung cancer, both through indirect effects on nicotine dependence and smoking behaviour, and through direct tumour cell-intrinsic effects [62–65]. A direct effect of *CHRNA5* expression on tumour cell-intrinsic growth, migration and invasion, has been supported by several *in vitro* studies, although the outcome is likely dependent on the expression pattern of other nAChR subunits expressed in each experimental system [66–69].

The regulated use of alternative splice donor sites within exon 5 generates several annotated *CHRNA5* isoforms that all use the canonical terminal exon 6 [70, 71] (Figure 7A). We have identified a novel transcript, referred to here as *CHRNA5[AluSz]*, which uses an intronic *AluSz* element as alternative terminal exon and polyadenylation site (Figure 7A). *AluSz* exonisation was confirmed by RT-PCR in HEK293T, lung adenocarcinoma A549 and esophageal adenocarcinoma OE19 cells (Figure S9), as well as by analysis of long-read RNA-seq data from HEK293T and A549 cells, and esophageal squamous cell carcinoma TE5 and normal immortalized esophageal squamous epithelial SHEE cells [72] (Figure S10A). Remarkably, *CHRNA5[AluSz]* appeared to be the dominant isoform in fully transformed A549 and TE5 cells, whereas the balance shifted in favour of the canonical isoform in HEK293T and non-transformed SHEE cells (Figure S10A). Predominant expression of the *CHRNA5[AluSz]* isoform was also apparent when assessed by RT-qPCR in A549 cells (Figure 7B), and was also observed in analysis of RNA-seq data from non-small cell lung carcinoma and esophageal squamous cell carcinoma cell lines in CCLE, whereas several neuroblastoma cell lines, originating from a tissue where *CHRNA5* is physiologically highly expressed, exhibited expression additionally of the canonical isoform (Figure S10B). These results implied that, despite not being previously identified, *CHRNA5[AluSz]* was the major isoform expressed particularly in cancer. Consistent, with this notion, both the canonical and the *CHRNA5[AluSz]* isoforms were highly upregulated in several cancer types, compared with the respective normal tissues, with expression of the *CHRNA5[AluSz]* approaching or often exceeding that of the canonical isoform (Figure S11).

**Figure 7.**
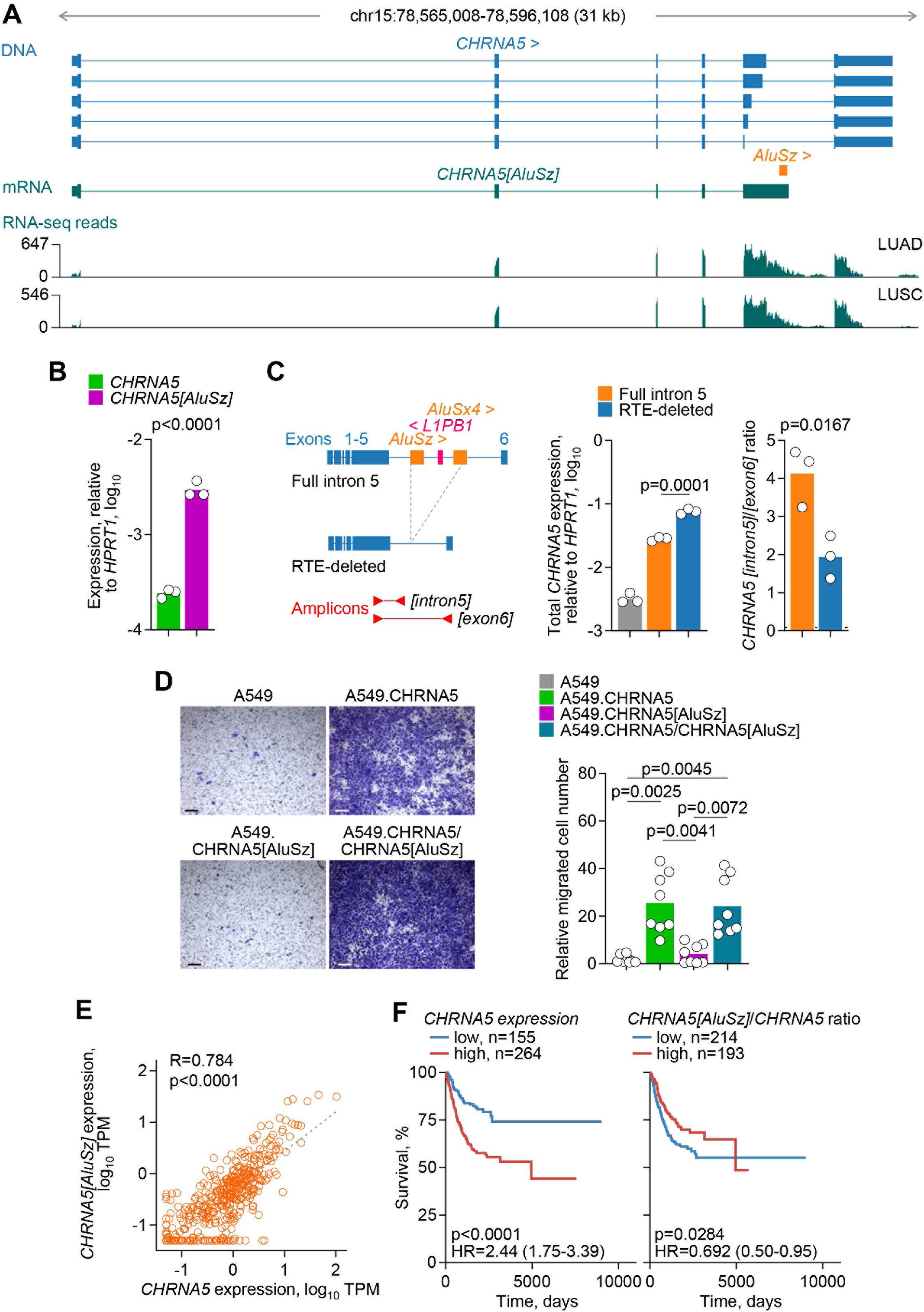
Characterisation of the *CHRNA5[AluSz]* isoform. **(A)** Gene structure and location of exonised *AluSz*, assembled *CHRNA5[AluSz]* transcript, and RNA-seq traces of 24 combined LUAD and LUSC samples at the *CHRNA5* locus. **(B)** *CHRNA5* and *CHRNA5[AluSz]* expression (assessed by RT-PCR and plotted relatively to *HPRT1* expression) in A549 cells. Symbols represent replicates (n=3) from a single experiment (p value calculated with two-tailed Student’s t-test). **(C)** *Left*, Schematic representation of *CHRNA5* cDNA minigene constructs retaining only intron 5 either with the reference complement of RTEs (full intron 5) or with the *AluSz* and adjacent *L1PB1* and *AluSx4* elements deleted (RTE-deleted), and of the amplicon used to measure expression. *Right*, total *CHRNA5* expression (assessed by RT-PCR and plotted relatively to *HPRT1* expression), and ratio of *CHRNA5[intron5]* to *CHRNA5[exon6]* in parental A549 cells or those transfected with either construct. Symbols represent replicates (n=3) from a single experiment (p values calculated with two-tailed Student’s t-tests between the full intron 5 and RTE-deleted transfections). **(D)** Representative crystal violet staining of *in vitro* migrated parental A549 cells and A549 cells expressing the canonical *CHRNA5*, the *CHRNA5[AluSz]* or both isoforms (*left*) (scale bar=200µm), and quantitation of the migrated cell number of each genotype (*right*). Symbols represent independent measurements (n=8, 4 fields of view from 2 independent experiments; p values calculated with Kruskal-Wallis test with Dunn’s multiple comparisons correction). **(E)** Correlation of *CHRNA5* and *CHRNA5[AluSz]* expression in LUAD samples (n=419) (p value calculated with linear regression). **(F)** Overall survival of LUAD patients, stratified by *CHRNA5* expression (*left*) or the ratio of *CHRNA5[AluSz]* to *CHRNA5* expression (*right*) (p values calculated with log-rank tests).

To examine the effect of the intronic RTEs on *CHRNA5[AluSz]* expression, we tested minigene constructs of *CHRNA5* cDNA retaining only intron 5 with the reference complement of RTEs or with the *AluSz* and adjacent *L1PB1* and *AluSx4* elements deleted (Figure 7C). Overall *CHRNA5* transcription from either minigene construct transfected into A549 cells exceeded endogenous CHRNA5 expression by at least an order of magnitude, but deletion of intronic RTEs resulted in higher levels of transcription compared with the full intron 5 (Figure 7C). Moreover, deletion of the intronic RTEs caused a significant shift in the balance of the two isoforms in favour of the canonical isoform terminating in exon 6 (*CHRNA5[exon6]*), although isoforms produced by continued transcription into intron 5 (*CHRNA5[intron5]*) still remained dominant (Figure 7C). These results suggested that, although not essential, the presence of the specific RTEs in *CHRNA5* intron 5 favour the production of the intronically-terminated *CHRNA5* isoform over the canonical.

Given its high expression, we next assessed the potential of the *CHRNA5[AluSz]* isoform to produce a functional protein. At the protein level, *AluSz* exonisation replaces the last 52 amino acids, which are encoded by canonical exon 6 and include the 4^th^ transmembrane helix of CHRNA5, with as shorter sequence, predicted to remain cytoplasmic (Figure S12A, B). Expression of influenza hemagglutinin (HA)-tagged versions of the canonical CHRNA5 and CHRNA5[AluSz] protein isoforms showed equivalent cell-surface expression in HEK293T, visualised by immunofluorescence (Figure S12C), indicating efficient translation and plasma membrane trafficking of both. To examine the potential biological activity of the CHRNA5[AluSz] protein we stably expressed either isoform in A549 cells (Figure S13). Neither isoform significantly affected the *in vitro* growth rate of A549 cells (Figure S14). Consistent with prior reports [68, 69], expression of the canonical isoform dramatically enhanced migration of A549 cells, whereas expression of CHRNA5[AluSz] had no apparent effect (Figure 7D). These results suggested that the loss of the last transmembrane helix resulted in a non-functional CHRNA5 isoform. Interestingly, acquired mutations resulting in an identical truncation of CHRNA6 have been found responsible for evolved insect resistance to insecticides [73], further highlighting the essential function of the last transmembrane helix. To test whether incorporation of the truncated CHRNA5[AluSz] protein into heteropentamers could potentially interfere with the function of the canonical, we co-expressed both isoforms in A549 cells (Figure S13). In this setting, cell migration was still significantly enhanced in doubly-expressing A549 cells, at levels comparable with those of A549 cells expressing the canonical isoform only (Figure 7D). These findings argued against a negative effect of CHRNA5[AluSz], in agreement with the lack of ligand binding by the CHRNA5 subunit [61].

Collectively, these results indicated that the switch to *CHRNA5[AluSz]* expression we observed in cancer would severely compromise the levels of canonical *CHRNA5* that would otherwise be produced and, in turn, reduce the pro-tumour effects of *CHRNA5* expression. A similar effect could also be achieved by alternative splicing within exon 5, causing a frame-shift in the translation of the exon 6 and producing a similarly truncated isoform (NCBI ID: NP_001382100). Although other alternative, non-functional isoforms can be detected [70, 71], the *CHRNA5[AluSz]* appears to be the dominant isoform (Figure S10). Expression of *CHRNA5[AluSz]* was significantly correlated with that of the canonical isoform in LUAD samples, although their ratio varied considerably among individual cases (Figure 7E). In agreement with a pro-tumour role, high overall *CHRNA5* transcription was associated with worse prognosis in LUAD (Figure 7F). However, a higher fraction of *CHRNA5* transcription diverted to the *CHRNA5[AluSz]* isoform was associated with better prognosis in the same cohort (Figure 7F), suggesting a protective effect of a switch to the non-functional isoform.

## Discussion

Evolutionary selection against deleterious effects is constantly depleting the germline of RTE integrations that pose a threat to nearby genes [74]. Nevertheless, numerous recent germline RTE integrations can adversely affect gene function, when the mechanisms that normally prevent their transcription and inclusion in gene transcripts fail. Our data indicate that transcriptional activation of RTEs causes widespread disruption of the transcriptional programme in cancer. Although they would be expected to be counterselected during tumour evolution, we identified several exemplar cases where transcriptional activation of embedded RTEs disrupts the function of a tumour-promoting or essential gene.

Transcriptionally activated RTEs appear to disrupt the function of adjacent protein-coding genes by two main mechanisms. The first is reduction in the transcription of the protein-coding isoform by RTE-initiated antisense transcription, as exemplified here by *RNGTT*, *CDH4*, *TLR7* and *APOBEC3B*. Antisense transcription has long been recognised as a mechanism of gene regulation more broadly [75]. Furthermore, RTEs have also been implicated in the initiation of *cis* antisense transcripts that may regulate gene expression under physiological conditions [76]. Of note, RTE integrations driving *cis* natural antisense transcripts are enriched near the 3’ UTR of genes and belong to relatively older *L2* and *MIR* subfamilies of non-LTR elements, implying they have been selected during evolution [76]. In contrast, RTEs identified here as regulators of cancer-promoting genes are primarily intergenic or intronic integrations of HERVs and other LTR elements, suggesting that the transcriptional activation of otherwise suppressed RTEs may extend regulation by antisense transcription to a new set of genes specifically in cancer.

The second mechanism by which transcriptionally activated RTEs can disrupt gene function is a switch to the production of non-functional isoforms by RTE-exonisation and alternative splicing. Switch to a non-functional isoform can be at the expense of the canonical protein-coding isoform, with mutually exclusive expression of the two. However, non-functional RTE-exonising isoforms may also be expressed proportionally with the canonical, yet considerably reduce the functional output the gene would otherwise produce. Such an effect on gene function would still be strong, particularly when the non-functional isoform becomes the dominant isoform, as in the case of *CHRNA5[AluSz]*. The switch to non-functional isoforms appears to involve younger RTEs of the *Alu* and *L1* subfamilies, the transcriptional utilisation of which is shared by diverse cancer types. A cancer-specific switch to non-functional isoforms may also explain the previously noted poor correlation between abundance of RNA transcripts, the quantitation of which often ignores the functional potential, and protein levels encoded from at least some genes in cancer [77].

Widespread RTE-mediated loss of function of tumour-promoting genes, as suggested by our findings, is seemingly at odds with the expected effect on tumour fitness that would disadvantage such events during tumour evolution, but may be further supported by recent evidence. A hypoxia-responsive *LTR12B* RTE has been reported to act as a cryptic promoter of an alternative isoform of *POU5F1*, encoding the pluripotency transcription factor OCT4, producing a likely non-functional version of this tumour-promoting protein in renal cell carcinoma [78]. Similarly, antisense transcription has been reported to regulate levels of the E3 ubiquitin ligase HECTD2, which would otherwise exert a clear tumour-promoting effect in melanoma [79].

Transcriptional activation of intronic RTEs has also been linked with incomplete mRNA splicing, which reduces levels of fully-spliced, functional mRNA isoforms and, consequently, tumour cell fitness [80]. Although prior examples in cancer may be limited, similar events have been reported to affect gene function also in physiological conditions. For example, a truncated, non-functional form of ACE2 is produced during infection or inflammation by an IFN-responsive *MIRb* element, acting as an alternative promoter [81]. The use of an intronic *L2a* element as an alternative terminal exon creates a *CD274* isoform that encodes a soluble version of PD-L1, which not only lacks suppressive activity, but also antagonises the membrane-bound canonical PD-L1 [82]. Similarly, the use of an intronic *Alu* element creates an isoform of *IFNAR2*, encoding a truncated version of the type I IFN receptor subunit 2, acting as a decoy receptor [83].

Collectively, these findings underscore the mutagenic potential of RTE insertions, which may be higher than previously appreciated and further enhanced in cancer by their release from epigenetic control. Dysregulation of RTEs in cancer is considered to serve as a warning signal for the emergence of transformed cells. Transcriptional activation of RTEs creates immunogenic ligands that are recognised by innate immune sensors and adaptive antigen receptors, thereby contributing to tumour immunogenicity and immune control [84, 85]. A potential effect of transcriptionally activated RTEs on the function of tumour-promoting or essential genes may represent an additional barrier to transformation. Similar to the immunogenic functions of transcriptionally activated RTEs, disruption of the cancer transcriptional programme would be subject to counterselection during evolution of individual tumours, but it may be positively selected during the evolution of the host species. Whereas the evolution of new function from RTE exaptation, particularly their utilisation in functional proteins is thought to be a slow evolutionary process [86], the regulation of adjacent gene function by co-option of transcriptionally metastable RTE integrations may evolve faster.

Several of the genes affected by transcriptional activation of RTEs are known to exert strong cell-intrinsic pro-tumour effects. Considered in isolation, this finding would support a potential anti-tumour role for RTE dysregulation through the disruption of the function of those genes. However, there are a number of confounding factors that increase the complexity of these effects.

Some of the affected genes are pleiotropic, with both tumour cell-intrinsic effects and effect on the immune or stroma microenvironment that can indirectly influence tumour grown. Direct indirect effects of an affected gene can synergise to promote tumour growth. For example, HECTD2 drives tumour cell-intrinsic proliferation of melanoma cells, as well as the production of immunosuppressive mediators [79], whereas ectopic expression of *CALB1* prevents senescence of squamous lung carcinoma cells, but also prevents pro-tumour recruitment of neutrophils by cytokines that would otherwise be secreted as part of the senescence-associated secretory phenotype [11]. Similarly, ENPP3 promotes cell-intrinsic growth and migration renal cell carcinoma cells [56, 60], but also regulates the availability of STING ligands for immune cells [59], and given its central role in RNA capping, *RNGTT* has the potential to affect many other genes with indirect effects on tumour growth.

Moreover, while the function of a gene may be clearly pro-tumour in the context of an established tumour or cell line, it may play a different role at a different stage during cancer initiation and progression. It may also be the case that the relative fitness cost incurred by RTE-mediated disruption of pro-tumour gene function is a late event in tumour progression, by which time clonal competition between tumour cells has taken place. It may also be that such fitness costs are an unavoidable consequence of global RTE activation during tumour evolution, but offset by gains in tumour-promoting functions resulting from the same underlying epigenetic changes, so that the net effect on tumour growth is positive, and the effect on each gene has to be considered in the context of all other changes. Lastly, an overall negative effect on tumour cell-intrinsic growth caused of RTE-mediated disruption of the cancer transcriptional programme may still benefit tumours by restraining the exponential growth of late-stage tumours that would otherwise outrun or outpace available resources.

Regardless of the ultimate effect on tumour growth, the identification and characterisation of specific cases of tumour-promoting genes affected by metastable RTE integrations highlights their potential to disrupt gene function in cancer, in turn increasing our understanding of tumour evolution and offering opportunities for intervention.

## Data availability

The RNA-seq data generated in this study have been deposited at the EMBL-EBI repository (www.ebi.ac.uk/arrayexpress) (E-MTAB-14514). TCGA and GTEx data used for the analyses described in this manuscript were obtained from dbGaP (https://dbgap.ncbi.nlm.nih.gov) accession numbers phs000178.v10.p8.c1 and phs000424.v7.p2.c1 in 2017. Other publicly available dataset supporting the findings of this study included the following: RNA-seq data from a renal cell carcinoma cell line RCC4 with restored expression of the Von Hippel-Lindau (VHL) tumour suppressor protein (GSE120887) [28]; ISO-seq data from ESCC cell line TE5 and normal immortalized esophageal squamous epithelial cell line SHEE (PRJNA515570) [72]; Long-read RNA-seq data from HEK293T and A549 cells (https://github.com/GoekeLab/sg-nex-data).

## Acknowledgements

We are grateful for assistance from the Advanced Sequencing, Cell Services, Flow cytometry, High Throughput Screening, Advanced Light Microscopy and Scientific Computing facilities at the Francis Crick Institute. The results shown here are in whole or part based upon data generated by the TCGA Research Network (http://cancergenome.nih.gov). The Genotype-Tissue Expression (GTEx) Project was supported by the Common Fund of the Office of the Director of the National Institutes of Health, and by NCI, NHGRI, NHLBI, NIDA, NIMH, and NINDS. This work was supported by the Francis Crick Institute (CC2088), which receives its core funding from Cancer Research UK, the UK Medical Research Council, and the Wellcome Trust. This project has received funding from the European Research Council (ERC) under the European Union’s Horizon 2020 research and innovation program (grant agreement No. 101018670). For the purpose of Open Access, the author has applied a CC BY public copyright license to any Author Accepted Manuscript version arising from this submission.

## References

1. Wells JN, Feschotte C: A Field Guide to Eukaryotic Transposable Elements. Annu Rev Genet 2020, 54:539–561.

2. Mills RE, Bennett EA, Iskow RC, Devine SE: Which transposable elements are active in the human genome? Trends Genet 2007, 23:183–191.

3. Fueyo R, Judd J, Feschotte C, Wysocka J: Roles of transposable elements in the regulation of mammalian transcription. Nat Rev Mol Cell Biol 2022, 23:481–497.

4. Modzelewski AJ, Gan Chong J, Wang T, He L: Mammalian genome innovation through transposon domestication. Nature Cell Biology 2022, 24:1332–1340.

5. Chuong EB, Elde NC, Feschotte C: Regulatory activities of transposable elements: from conflicts to benefits. Nat Rev Genet 2017, 18:71–86.

6. Ishak CA, De Carvalho DD: Reactivation of Endogenous Retroelements in Cancer Development and Therapy. Annu Rev Cancer Biol 2020, 4:159–176.

7. Burns KH: Transposable elements in cancer. Nat Rev Cancer 2017, 17:415–424.

8. Attig J, Young GR, Hosie L, Perkins D, Encheva-Yokoya V, Stoye JP, Snijders AP, Ternette N, Kassiotis G: LTR retroelement expansion of the human cancer transcriptome and immunopeptidome revealed by de novo transcript assembly. Genome Res 2019, 29:1578–1590.

9. Lamprecht B, Walter K, Kreher S, Kumar R, Hummel M, Lenze D, Köchert K, Bouhlel MA, Richter J, Soler E, et al: Derepression of an endogenous long terminal repeat activates the CSF1R proto-oncogene in human lymphoma. Nat Med 2010, 16:571–579, 571p following 579.

10. Babaian A, Romanish MT, Gagnier L, Kuo LY, Karimi MM, Steidl C, Mager DL: Onco-exaptation of an endogenous retroviral LTR drives IRF5 expression in Hodgkin lymphoma. Oncogene 2016, 35:2542–2546.

11. Attig J, Pape J, Doglio L, Kazachenka A, Ottina E, Young GR, Enfield KS, Aramburu IV, Ng KW, Faulkner N, et al: Human endogenous retrovirus onco-exaptation counters cancer cell senescence through Calbindin. J Clin Invest 2023.

12. Wiesner T, Lee W, Obenauf AC, Ran L, Murali R, Zhang QF, Wong EW, Hu W, Scott SN, Shah RH, et al: Alternative transcription initiation leads to expression of a novel ALK isoform in cancer. Nature 2015, 526:453–457.

13. Jang HS, Shah NM, Du AY, Dailey ZZ, Pehrsson EC, Godoy PM, Zhang D, Li D, Xing X, Kim S, et al: Transposable elements drive widespread expression of oncogenes in human cancers. Nat Genet 2019, 51:611–617.

14. Tange O: GNU Parallel: The Command-Line Power Tool. The USENIX Magazine 2011, 36:42–47.

15. Danecek P, Bonfield JK, Liddle J, Marshall J, Ohan V, Pollard MO, Whitwham A, Keane T, McCarthy SA, Davies RM, Li H: Twelve years of SAMtools and BCFtools. Gigascience 2021, 10.

16. Patro R, Duggal G, Love MI, Irizarry RA, Kingsford C: Salmon provides fast and bias-aware quantification of transcript expression. Nat Methods 2017, 14:417–419.

17. Thorvaldsdóttir H, Robinson JT, Mesirov JP: Integrative Genomics Viewer (IGV): high-performance genomics data visualization and exploration. Brief Bioinform 2013, 14:178–192.

18. Li H: Minimap2: pairwise alignment for nucleotide sequences. Bioinformatics 2018, 34:3094–3100.

19. Tang AD, Soulette CM, van Baren MJ, Hart K, Hrabeta-Robinson E, Wu CJ, Brooks AN: Full-length transcript characterization of SF3B1 mutation in chronic lymphocytic leukemia reveals downregulation of retained introns. Nat Commun 2020, 11:1438.

20. Lombardi O, Li R, Halim S, Choudhry H, Ratcliffe PJ, Mole DR: Pan-cancer analysis of tissue and single-cell HIF-pathway activation using a conserved gene signature. Cell Rep 2022, 41:111652.

21. Attig J, Young GR, Stoye JP, Kassiotis G: Physiological and Pathological Transcriptional Activation of Endogenous Retroelements Assessed by RNA-Sequencing of B Lymphocytes. Front Microbiol 2017, 8:2489.

22. Raudvere U, Kolberg L, Kuzmin I, Arak T, Adler P, Peterson H, Vilo J: g:Profiler: a web server for functional enrichment analysis and conversions of gene lists (2019 update). Nucleic Acids Res 2019, 47:191–198.

23. Tsherniak A, Vazquez F, Montgomery PG, Weir BA, Kryukov G, Cowley GS, Gill S, Harrington WF, Pantel S, Krill-Burger JM, et al: Defining a Cancer Dependency Map. Cell 2017, 170:564–576.e516.

24. Takahashi Y, Harashima N, Kajigaya S, Yokoyama H, Cherkasova E, McCoy JP, Hanada K, Mena O, Kurlander R, Tawab A, et al: Regression of human kidney cancer following allogeneic stem cell transplantation is associated with recognition of an HERV-E antigen by T cells. J Clin Invest 2008, 118:1099–1109.

25. Cherkasova E, Malinzak E, Rao S, Takahashi Y, Senchenko VN, Kudryavtseva AV, Nickerson ML, Merino M, Hong JA, Schrump DS, et al: Inactivation of the von Hippel-Lindau tumor suppressor leads to selective expression of a human endogenous retrovirus in kidney cancer. Oncogene 2011, 30:4697–4706.

26. Smith CC, Beckermann KE, Bortone DS, De Cubas AA, Bixby LM, Lee SJ, Panda A, Ganesan S, Bhanot G, Wallen EM, et al: Endogenous retroviral signatures predict immunotherapy response in clear cell renal cell carcinoma. J Clin Invest 2018, 128:4804–4820.

27. Au L, Hatipoglu E, Robert de Massy M, Litchfield K, Beattie G, Rowan A, Schnidrig D, Thompson R, Byrne F, Horswell S, et al: Determinants of anti-PD-1 response and resistance in clear cell renal cell carcinoma. Cancer Cell 2021, 39:1497–1518.e1411.

28. Smythies JA, Sun M, Masson N, Salama R, Simpson PD, Murray E, Neumann V, Cockman ME, Choudhry H, Ratcliffe PJ, Mole DR: Inherent DNA-binding specificities of the HIF-1α and HIF-2α transcription factors in chromatin. EMBO Rep 2019, 20.

29. Borden K, Culjkovic-Kraljacic B, Cowling VH: To cap it all off, again: dynamic capping and recapping of coding and non-coding RNAs to control transcript fate and biological activity. Cell Cycle 2021, 20:1347–1360.

30. Borden KLB: Cancer cells hijack RNA processing to rewrite the message. Biochem Soc Trans 2022, 50:1447–1456.

31. Faraji F, Hu Y, Wu G, Goldberger NE, Walker RC, Zhang J, Hunter KW: An integrated systems genetics screen reveals the transcriptional structure of inherited predisposition to metastatic disease. Genome Res 2014, 24:227–240.

32. Inuzuka H, Miyatani S, Takeichil M: R-cadherin: a novel Ca(2+)-dependent cell-cell adhesion molecule expressed in the retina. Neuron 1991, 7:69–79.

33. Leckband DE, de Rooij J: Cadherin adhesion and mechanotransduction. Annu Rev Cell Dev Biol 2014, 30:291–315.

34. Kaszak I, Witkowska-Piłaszewicz O, Niewiadomska Z, Dworecka-Kaszak B, Ngosa Toka F, Jurka P: Role of Cadherins in Cancer-A Review. Int J Mol Sci 2020, 21.

35. Wheelock MJ, Shintani Y, Maeda M, Fukumoto Y, Johnson KR: Cadherin switching. J Cell Sci 2008, 121:727–735.

36. Haass NK, Smalley KS, Li L, Herlyn M: Adhesion, migration and communication in melanocytes and melanoma. Pigment Cell Res 2005, 18:150–159.

37. Kreizenbeck GM, Berger AJ, Subtil A, Rimm DL, Gould Rothberg BE: Prognostic significance of cadherin-based adhesion molecules in cutaneous malignant melanoma. Cancer Epidemiol Biomarkers Prev 2008, 17:949–958.

38. Maeda M, Johnson E, Mandal SH, Lawson KR, Keim SA, Svoboda RA, Caplan S, Wahl JK, 3rd, Wheelock MJ, Johnson KR: Expression of inappropriate cadherins by epithelial tumor cells promotes endocytosis and degradation of E-cadherin via competition for p120(ctn). Oncogene 2006, 25:4595–4604.

39. Axberg I, Ramstedt U, Patarroyo M, Beatty P, Wigzell H: Inhibition of natural killer cell cytotoxicity by a monoclonal antibody directed against adhesion-mediating protein gp 90 (CD18). Scand J Immunol 1987, 26:547–554.

40. Appolloni I, Barilari M, Caviglia S, Gambini E, Reisoli E, Malatesta P: A cadherin switch underlies malignancy in high-grade gliomas. Oncogene 2015, 34:1991–2002.

41. Miotto E, Sabbioni S, Veronese A, Calin GA, Gullini S, Liboni A, Gramantieri L, Bolondi L, Ferrazzi E, Gafà R, et al: Frequent aberrant methylation of the CDH4 gene promoter in human colorectal and gastric cancer. Cancer Res 2004, 64:8156–8159.

42. Lind NA, Rael VE, Pestal K, Liu B, Barton GM: Regulation of the nucleic acid-sensing Toll-like receptors. Nat Rev Immunol 2022, 22:224–235.

43. Vinuesa CG, Grenov A, Kassiotis G: Innate virus-sensing pathways in B cell systemic autoimmunity. Science 2023, 380:478–484.

44. Kaczanowska S, Joseph AM, Davila E: TLR agonists: our best frenemy in cancer immunotherapy. J Leukoc Biol 2013, 93:847–863.

45. Dajon M, Iribarren K, Cremer I: Dual roles of TLR7 in the lung cancer microenvironment. Oncoimmunology 2015, 4:e991615.

46. Cherfils-Vicini J, Platonova S, Gillard M, Laurans L, Validire P, Caliandro R, Magdeleinat P, Mami-Chouaib F, Dieu-Nosjean MC, Fridman WH, et al: Triggering of TLR7 and TLR8 expressed by human lung cancer cells induces cell survival and chemoresistance. J Clin Invest 2010, 120:1285–1297.

47. Grimmig T, Matthes N, Hoeland K, Tripathi S, Chandraker A, Grimm M, Moench R, Moll EM, Friess H, Tsaur I, et al: TLR7 and TLR8 expression increases tumor cell proliferation and promotes chemoresistance in human pancreatic cancer. Int J Oncol 2015, 47:857–866.

48. Chatterjee S, Crozet L, Damotte D, Iribarren K, Schramm C, Alifano M, Lupo A, Cherfils-Vicini J, Goc J, Katsahian S, et al: TLR7 promotes tumor progression, chemotherapy resistance, and poor clinical outcomes in non-small cell lung cancer. Cancer Res 2014, 74:5008–5018.

49. Dajon M, Iribarren K, Petitprez F, Marmier S, Lupo A, Gillard M, Ouakrim H, Victor N, Vincenzo DB, Joubert PE, et al: Toll like receptor 7 expressed by malignant cells promotes tumor progression and metastasis through the recruitment of myeloid derived suppressor cells. Oncoimmunology 2019, 8:e1505174.

50. Baglivo S, Bianconi F, Metro G, Gili A, Tofanetti FR, Bellezza G, Ricciuti B, Mandarano M, Teti V, Siggillino A, et al: Higher TLR7 Gene Expression Predicts Poor Clinical Outcome in Advanced NSCLC Patients Treated with Immunotherapy. Genes (Basel) 2021, 12.

51. Stavrou S, Ross SR: APOBEC3 Proteins in Viral Immunity. J Immunol 2015, 195:4565–4570.

52. Swanton C, McGranahan N, Starrett GJ, Harris RS: APOBEC Enzymes: Mutagenic Fuel for Cancer Evolution and Heterogeneity. Cancer Discov 2015, 5:704–712.

53. Petljak M, Dananberg A, Chu K, Bergstrom EN, Striepen J, von Morgen P, Chen Y, Shah H, Sale JE, Alexandrov LB, et al: Mechanisms of APOBEC3 mutagenesis in human cancer cells. Nature 2022, 607:799–807.

54. Durfee C, Temiz NA, Levin-Klein R, Argyris PP, Alsøe L, Carracedo S, Alonso de la Vega A, Proehl J, Holzhauer AM, Seeman ZJ, et al: Human APOBEC3B promotes tumor development in vivo including signature mutations and metastases. Cell Rep Med 2023, 4:101211.

55. Doñate F, Raitano A, Morrison K, An Z, Capo L, Aviña H, Karki S, Morrison K, Yang P, Ou J, et al: AGS16F Is a Novel Antibody Drug Conjugate Directed against ENPP3 for the Treatment of Renal Cell Carcinoma. Clin Cancer Res 2016, 22:1989–1999.

56. Von Roemeling CA, Marlow LA, Radisky DC, Rohl A, Larsen HE, Wei J, Sasinowska H, Zhu H, Drake R, Sasinowski M, et al: Functional genomics identifies novel genes essential for clear cell renal cell carcinoma tumor cell proliferation and migration. Oncotarget 2014, 5:5320–5334.

57. Borza R, Salgado-Polo F, Moolenaar WH, Perrakis A: Structure and function of the ecto-nucleotide pyrophosphatase/phosphodiesterase (ENPP) family: Tidying up diversity. J Biol Chem 2022, 298:101526.

58. Tsai SH, Kinoshita M, Kusu T, Kayama H, Okumura R, Ikeda K, Shimada Y, Takeda A, Yoshikawa S, Obata-Ninomiya K, et al: The ectoenzyme E-NPP3 negatively regulates ATP-dependent chronic allergic responses by basophils and mast cells. Immunity 2015, 42:279–293.

59. Mardjuki R, Wang S, Carozza J, Zirak B, Subramanyam V, Abhiraman G, Lyu X, Goodarzi H, Li L: Identification of the extracellular membrane protein ENPP3 as a major cGAMP hydrolase and innate immune checkpoint. Cell Rep 2024, 43:114209.

60. Yano Y, Hayashi Y, Sano K, Nagano H, Nakaji M, Seo Y, Ninomiya T, Yoon S, Yokozaki H, Kasuga M: Expression and localization of ecto-nucleotide pyrophosphatase/phosphodiesterase I-1 (E-NPP1/PC-1) and −3 (E-NPP3/CD203c/PD-Ibeta/B10/gp130(RB13-6)) in inflammatory and neoplastic bile duct diseases. Cancer Lett 2004, 207:139–147.

61. Improgo MR, Scofield MD, Tapper AR, Gardner PD: The nicotinic acetylcholine receptor CHRNA5/A3/B4 gene cluster: dual role in nicotine addiction and lung cancer. Prog Neurobiol 2010, 92:212–226.

62. Amos CI, Wu X, Broderick P, Gorlov IP, Gu J, Eisen T, Dong Q, Zhang Q, Gu X, Vijayakrishnan J, et al: Genome-wide association scan of tag SNPs identifies a susceptibility locus for lung cancer at 15q25.1. Nat Genet 2008, 40:616–622.

63. Hung RJ, McKay JD, Gaborieau V, Boffetta P, Hashibe M, Zaridze D, Mukeria A, Szeszenia-Dabrowska N, Lissowska J, Rudnai P, et al: A susceptibility locus for lung cancer maps to nicotinic acetylcholine receptor subunit genes on 15q25. Nature 2008, 452:633–637.

64. Spitz MR, Amos CI, Dong Q, Lin J, Wu X: The CHRNA5-A3 region on chromosome 15q24-25.1 is a risk factor both for nicotine dependence and for lung cancer. J Natl Cancer Inst 2008, 100:1552–1556.

65. Thorgeirsson TE, Geller F, Sulem P, Rafnar T, Wiste A, Magnusson KP, Manolescu A, Thorleifsson G, Stefansson H, Ingason A, et al: A variant associated with nicotine dependence, lung cancer and peripheral arterial disease. Nature 2008, 452:638–642.

66. Krais AM, Hautefeuille AH, Cros MP, Krutovskikh V, Tournier JM, Birembaut P, Thépot A, Paliwal A, Herceg Z, Boffetta P, et al: CHRNA5 as negative regulator of nicotine signaling in normal and cancer bronchial cells: effects on motility, migration and p63 expression. Carcinogenesis 2011, 32:1388–1395.

67. Improgo MR, Soll LG, Tapper AR, Gardner PD: Nicotinic acetylcholine receptors mediate lung cancer growth. Front Physiol 2013, 4:251.

68. Chen X, Jia Y, Zhang Y, Zhou D, Sun H, Ma X: α5-nAChR contributes to epithelial-mesenchymal transition and metastasis by regulating Jab1/Csn5 signalling in lung cancer. J Cell Mol Med 2020, 24:2497–2506.

69. Wang ML, Hsu YF, Liu CH, Kuo YL, Chen YC, Yeh YC, Ho HL, Wu YC, Chou TY, Wu CW: Low-Dose Nicotine Activates EGFR Signaling via α5-nAChR and Promotes Lung Adenocarcinoma Progression. Int J Mol Sci 2020, 21.

70. Warzecha CC, Shen S, Xing Y, Carstens RP: The epithelial splicing factors ESRP1 and ESRP2 positively and negatively regulate diverse types of alternative splicing events. RNA Biol 2009, 6:546–562.

71. Falvella FS, Alberio T, Noci S, Santambrogio L, Nosotti M, Incarbone M, Pastorino U, Fasano M, Dragani TA: Multiple isoforms and differential allelic expression of CHRNA5 in lung tissue and lung adenocarcinoma. Carcinogenesis 2013, 34:1281–1285.

72. Cheng YW, Chen YM, Zhao QQ, Zhao X, Wu YR, Chen DZ, Liao LD, Chen Y, Yang Q, Xu LY, et al: Long Read Single-Molecule Real-Time Sequencing Elucidates Transcriptome-Wide Heterogeneity and Complexity in Esophageal Squamous Cells. Front Genet 2019, 10:915.

73. Baxter SW, Chen M, Dawson A, Zhao JZ, Vogel H, Shelton AM, Heckel DG, Jiggins CD: Mis-spliced transcripts of nicotinic acetylcholine receptor alpha6 are associated with field evolved spinosad resistance in Plutella xylostella (L.). PLoS Genet 2010, 6:e1000802.

74. Sultana T, Zamborlini A, Cristofari G, Lesage P: Integration site selection by retroviruses and transposable elements in eukaryotes. Nat Rev Genet 2017, 18:292–308.

75. Pelechano V, Steinmetz LM: Gene regulation by antisense transcription. Nat Rev Genet 2013, 14:880–893.

76. Conley AB, Miller WJ, Jordan IK: Human cis natural antisense transcripts initiated by transposable elements. Trends Genet 2008, 24:53–56.

77. Zhang B, Wang J, Wang X, Zhu J, Liu Q, Shi Z, Chambers MC, Zimmerman LJ, Shaddox KF, Kim S, et al: Proteogenomic characterization of human colon and rectal cancer. Nature 2014, 513:382–387.

78. Siebenthall KT, Miller CP, Vierstra JD, Mathieu J, Tretiakova M, Reynolds A, Sandstrom R, Rynes E, Haugen E, Johnson A, et al: Integrated epigenomic profiling reveals endogenous retrovirus reactivation in renal cell carcinoma. EBioMedicine 2019, 41:427–442.

79. Ottina E, Panova V, Doglio L, Kazachenka A, Cornish G, Kirkpatrick J, Attig J, Young GR, Litchfield K, Lesluyes T, et al: E3 ubiquitin ligase HECTD2 mediates melanoma progression and immune evasion. Oncogene 2021, 40:5567–5578.

80. Kazachenka A, Loong JH, Attig J, Young GR, Ganguli P, Devonshire G, Grehan N, Ciccarelli FD, Fitzgerald RC, Kassiotis G: The transcriptional landscape of endogenous retroelements delineates esophageal adenocarcinoma subtypes. NAR Cancer 2023, 5:zcad040.

81. Ng KW, Attig J, Bolland W, Young GR, Major J, Wrobel AG, Gamblin S, Wack A, Kassiotis G: Tissue-specific and interferon-inducible expression of nonfunctional ACE2 through endogenous retroelement co-option. Nat Genet 2020, 52:1294–1302.

82. Ng KW, Attig J, Young GR, Ottina E, Papamichos SI, Kotsianidis I, Kassiotis G: Soluble PD-L1 generated by endogenous retroelement exaptation is a receptor antagonist. Elife 2019, 8.

83. Pasquesi GIM, Allen H, Ivancevic A, Barbachano-Guerrero A, Joyner O, Guo K, Simpson DM, Gapin K, Horton I, Nguyen L, et al: Regulation of human interferon signaling by transposon exonization. bioRxiv 2023.

84. Lindholm HT, Chen R, De Carvalho DD: Endogenous retroelements as alarms for disruptions to cellular homeostasis. Trends Cancer 2023, 9:55–68.

85. Kassiotis G: The Immunological Conundrum of Endogenous Retroelements. Annu Rev Immunol 2023, 41:99–125.

86. Gotea V, Makałowski W: Do transposable elements really contribute to proteomes? Trends Genet 2006, 22:260–267.

