## Supplemental_Data for "Retroelement co-option disrupts the cancer transcriptional programme"

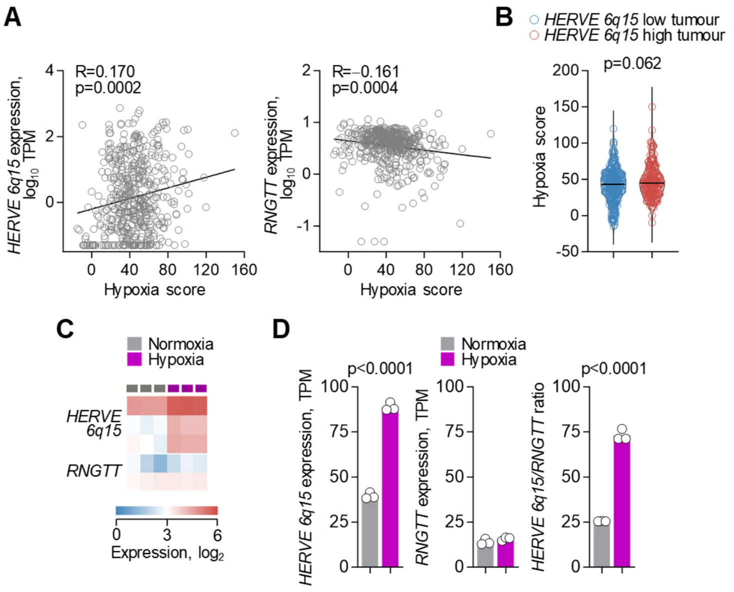


**Figure S1.** ***HERVE 6q15* responsiveness to hypoxia. (A)** *HERVE 6q15* and *RNGTT* expression (TPM) in KIRC samples (n=485) according to their hypoxia scores (p values calculated with linear regression). **(B)** Hypoxia score in KIRC samples with low (n=285) and high (n=200) *HERVE 6q15* expression (p value calculated with Mann-Whitney test). **(C)** Expression of transcripts overlapping *HERVE 6q15* or the canonical *RNGTT* in RNA-seq data (GSE120887) from VHL-sufficient RCC4 cells grown in normoxic or hypoxic conditions. **(D)** *HERVE 6q15* and *RNGTT* expression (TPM), and ratio of *HERVE 6q15* to *RNGTT* expression in the same cells as in c (p values calculated with two-tailed Student's t-tests).

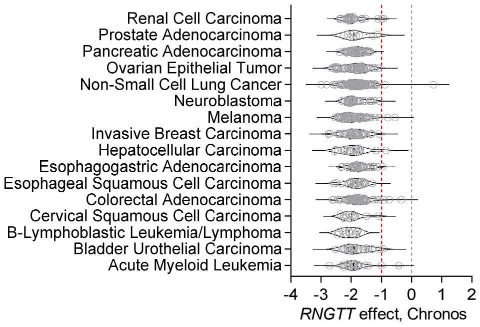


**Figure S2. Essential function of *RNGTT* in cancer cell lines.** Effect of *RNGTT* deletion (Chronos score) on the growth of cancer cell lines of the indicated origin. Red dashed line represents the threshold for a significant effect. Data downloaded from the Dependency Map (DepMap) portal (https://depmap.org/portal).

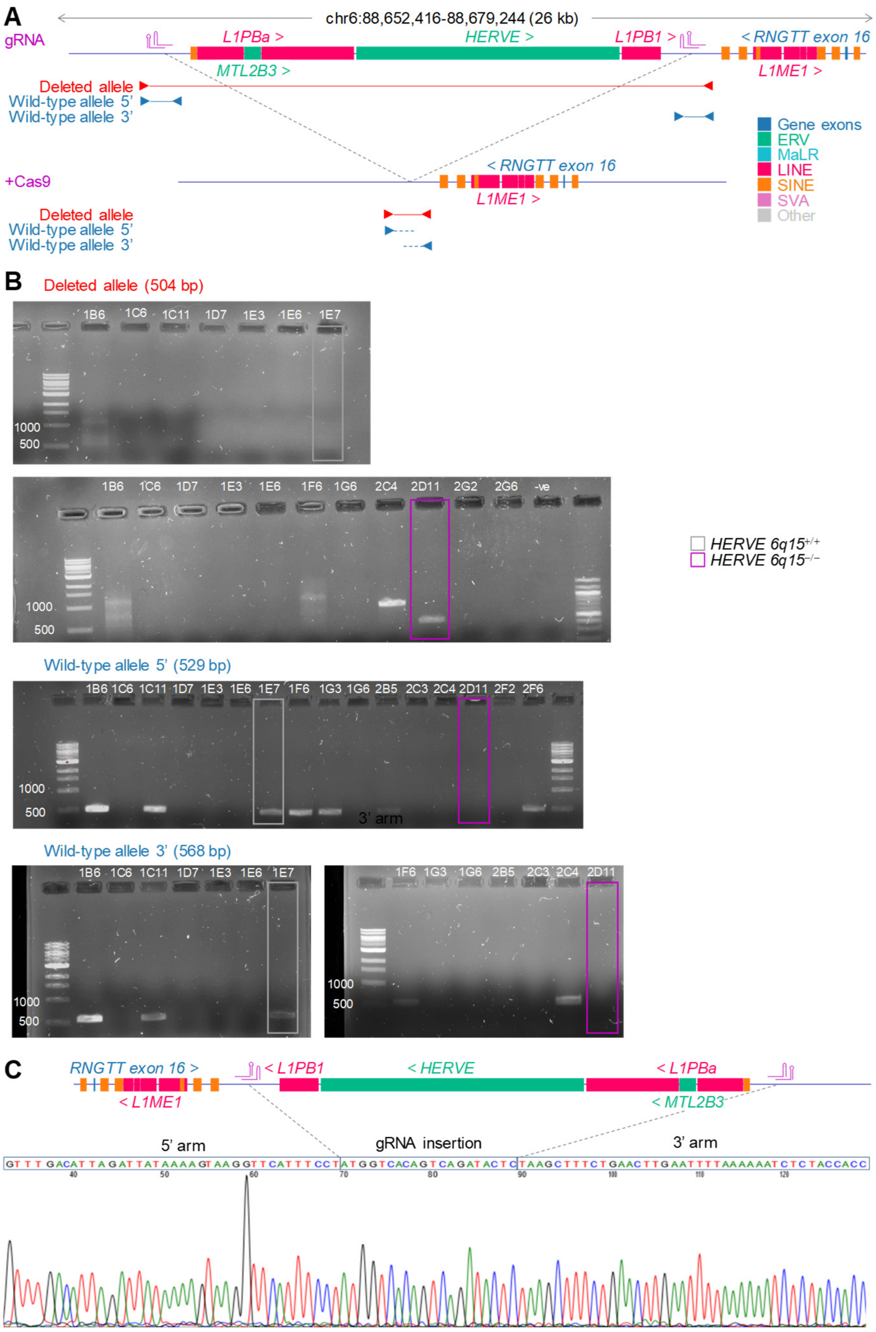


**Figure S3. CRISPR/Cas9-mediated deletion of the *HERVE 6q15* provirus. (A)** RTE content in the intronic region of *HERVE 6q15* integration, position of guide RNA (gRNA) molecules, and position of the PCR primers used for the detection of the wild-type and deleted alleles. **(B)** Gel electrophoresis of amplicons from A498 clones with wild-type or deleted alleles. **(C)** Sanger sequencing of PCR amplicon of the deleted allele in *HERVE 6q15*$^{-/-}$ clone 2D11. Also noted is the insertion of one gRNA sequence in site of the deletion.

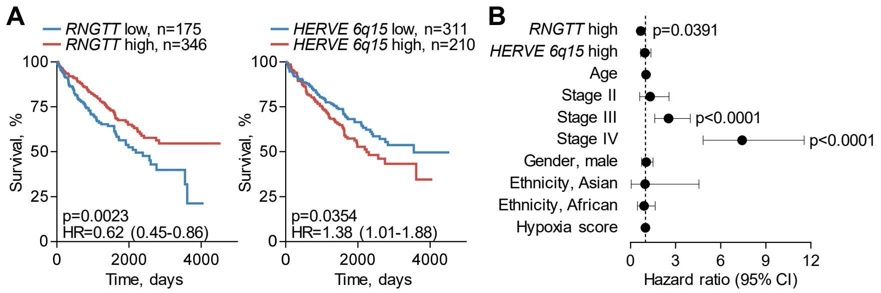


**Figure S4. Correlation of *RNGTT* and *HERVE 6q15* expression with survival in KIRC. (A)** Overall survival of KIRC patients, stratified by expression levels of *RNGTT* (*left*) or *HERVE 6q15* (*right*) (p values calculated with log-rank tests). **(B)** Overall survival hazard ratios (HRs) for the indicated variables in KIRC patients (*RNGTT* high n=346, reference low n=175; *HERVE 6q15* high n=210, reference low n=311; Age n=521; Stage II n=56, III n=123, IV n=82, reference I n=257; Gender, male n=338, reference female n=183; Ethnicity, Asian n=8, African n=55, reference White n=450; Hypoxia score n=470). Error bars represent 95% confidence intervals (CIs) (p values calculated with Cox proportional hazards regression).

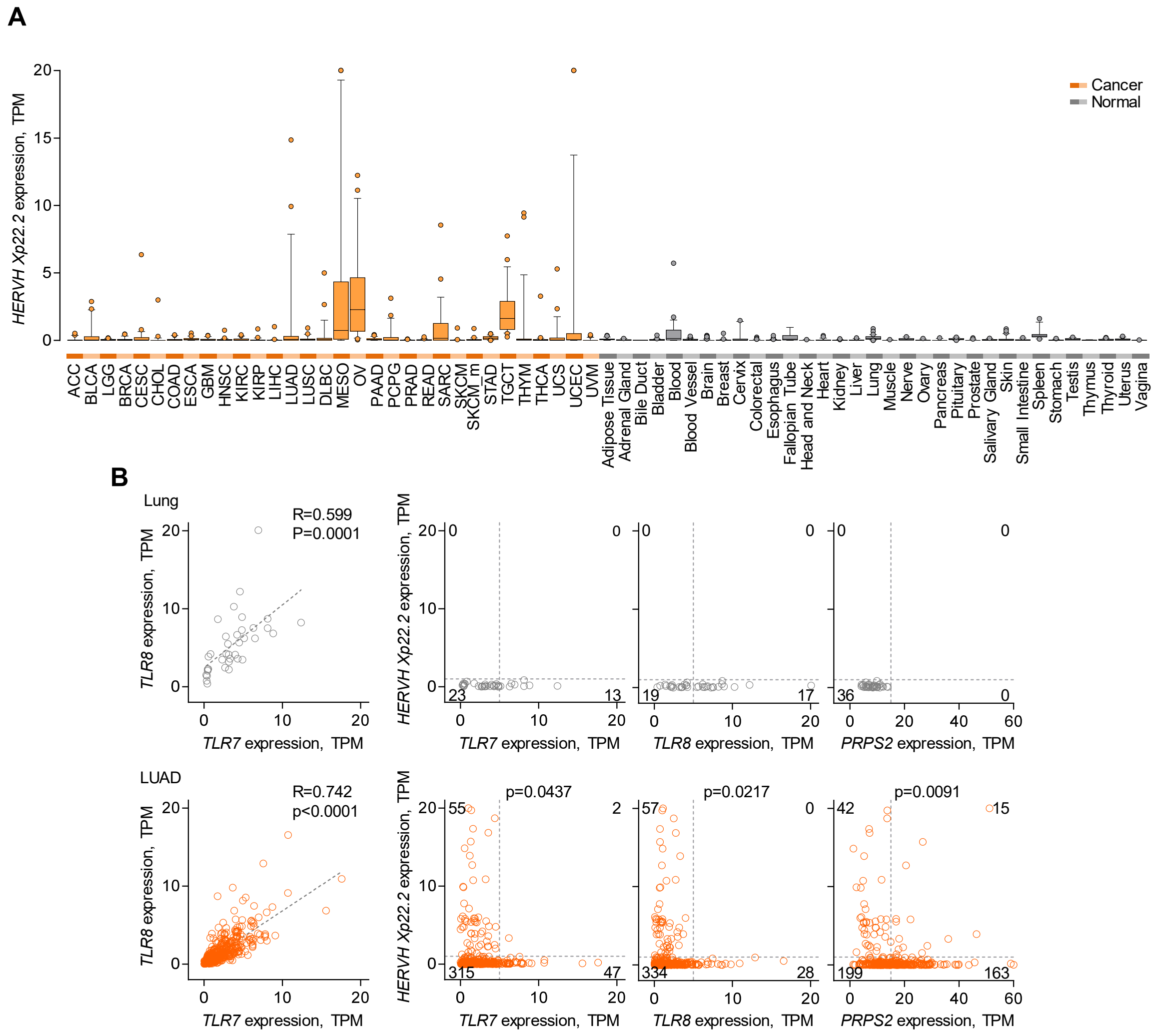


**Figure S5. *HERVH Xp22.2* expression across cancers and correlation with *TLR7*, *TLR8* and *PRPS2* expression in LUAD. (A)** *HERVH Xp22.2* expression (TPM) in the indicated TCGA cancer types (n = 24 per type) and in normal tissues from GTEx (n = 2-156 per tissue type). Box plots denote median value and quartiles, whiskers denote 1.5x the interquartile range, and individual points denote outliers. Values are capped at 20 TPM. **(B)** Correlation of *HERVH Xp22.2*, *TLR7*, *TLR8* and *PRPS2* expression (TPM) in normal lung tissue (n=36) or LUAD samples (n=419) (p values calculated with linear regression for positive correlations, and with Fisher's exact tests for negative correlations using the numbers in the quadrants).

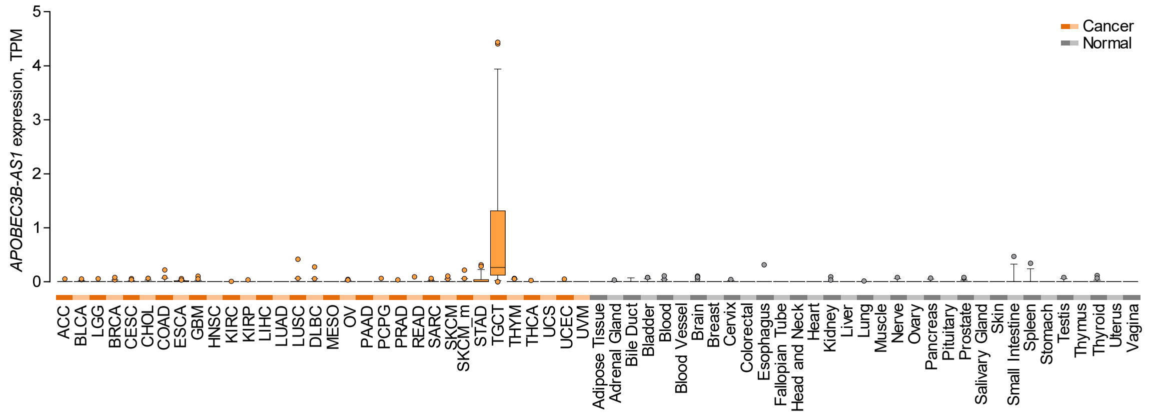


**Figure S6. *APOBEC3B-AS1* expression across cancers and normal tissues.** *APOBEC3B-AS1* expression (TPM) in the indicated TCGA cancer types (n = 24 per type) and in normal tissues from GTEx (n = 2-156 per tissue type). Box plots denote median value and quartiles, whiskers denote 1.5x the interquartile range, and individual points denote outliers.

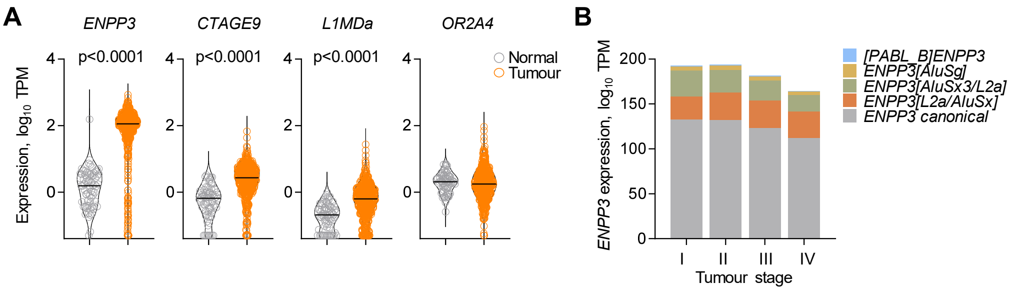


**Figure S7. Cancer specificity of *ENNP3*, and overlapping gene and RTE expression in KIRC. (A)** Expression of canonical *ENNP3* or overlapping genes and RTEs in normal kidney tissue (n=71) and KIRC samples (n=538) (p values calculated with Mann-Whitney tests). **(B)** *ENPP3* isoform expression (TPM) in KIRC samples according to tumour stage (I n=271, II n=59, III n=123, IV n=82).

**A**

```
                  1                                                                                                                      120
                  |        Transmembrane helix                                                                                             |
           ENPP3  MESTLTLATEQPVKKNTLKKYKIACIVLLALLVIMSLGLGLGLGLRKLEKQGSCRKKCFDASFRGLENCRCDVACKDRGDCCWDFEDTCVESTRIWMCNKFRCGETRLEASLCSCSDDCL
ENPP3[L2a/AluSx]  MESTLTLATEQPVKKNTLKKYKIACIVLLALLVIMSLGLGLGLGLRKLEKQGSCRKKCFDASFRGLENCRCDVACKDRGDCCWDFEDTCVESTRIWMCNKFRCGETRLEASLCSCSDDCL
                121                                                                                                                      240
                  |                                                                              Phosphodiesterase                         |
           ENPP3  QRKDCCADYKSVCQGETSWLEENCDTAQQSQCPEGFDLPPVILFSMDGFRAEYLYTWDTLMPNINKLKTCGIHSKYMRAMYPTKTFPNHYTIVTGLYPESHGIIDNNMYDVNLNKNFSLS
ENPP3[L2a/AluSx]  QRKDCCADYKSVCQGETSWLEENCDTAQQSQCPEGFDLPPVILFSMDGFRAEYLYTWDTLMPNINKLKTCGIHSKYMRAMYPTKTFPNHYTIVTGLYPESHGIIDNNMYDVNLNKNFSLS
                241                                                                                                                      360
                  |                                                                                                                        |
           ENPP3  SKEQNNPAWWHGQPMWLTAMYQGLKAATYFWPGSEVAINGSFPSIYMPYNGSVPFEERISTLLKWLDLPKAERPRFYTMYFEEPDSSGHAGGPVSARVIKALQVVDHAFGMLMEGLKQRN
ENPP3[L2a/AluSx]  SKEQNNPAWWHGQPMWLTAMYQGLKAATYFWPGSEVAINGSFPSIYMPYNGSVPFEERISTLLKWLDLPKAERPRFYTMYFEEPDSSGHAGGPVSARVIKALQVVDHAFGMLMEGLKQRN
                361                                                                                                                      480
                  |                                                                                                                        |
           ENPP3  LHNCVNIILLADHGMDQTYCNKMEYMTDYFPRINFFYMYEGPAPRIRAHNIPHDFFSFNSEEIVRNLSCRKPDQHFKPYLTPDLPKRLHYAKNVRIDKVHLFVDQQWLAVRSKSNTNCGG
ENPP3[L2a/AluSx]  LHNCVNIILLADHGMDQTYCNKMEYMTDYFPRINFFYMYEGPAPRIRAHNIPHDFFSFNSEEIVRNLSCRKPDQHFKPYLTPDLPKRLHYAKNVRIDKVHLFVDQQWLAVRFVYLFTYLI
                481                                                                                                                      600
                  |                                                                                                                        |
           ENPP3  GNHGYNNEFRSMEAIFLAHGPSFKEKTEVEPFENIEVYNLMCDLLRIQPAPNNGTHGSLNHLLKVPFYEPSHAEEVSKFSVCGFANPLPTESLDCFCPHLQNSTQLEQVNQMLNLTQEEI
ENPP3[L2a/AluSx]  MPLITICHYKST------------------------------------------------------------------------------------------------------------
                601                                                                                                                      720
                  |                                                                               Nuclease                                 |
           ENPP3  TATVKVNLPFGRPRVLQKNVDHCLLYHREYVSGFGKAMRMPMWSSYTVPQLGDTSPLPPTVPDCLRADVRVPPSESQKCSFYLADKNITHGFLYPPASNRTSDSQYDALITSNLVPMYEE
ENPP3[L2a/AluSx]  ------------------------------------------------------------------------------------------------------------------------
                720                                                                                                                      840
                  |                                                                                                                        |
           ENPP3  FRKMWDYFHSVLLIKHATERNGVNVVSGPIFDYNYDGHFDAPDEITKHLANTDVPIPTHYFVVLTSCKNKSHTPENCPGWLDVLPFIIPHRPTNVESCPEGKPEALWVEERFTAHIARVR
ENPP3[L2a/AluSx]  ------------------------------------------------------------------------------------------------------------------------
                841                               875
                  |                                 |
           ENPP3  DVELLTGLDFYQDKVQPVSEILQLKTYLPTFETTI
ENPP3[L2a/AluSx]  -----------------------------------
```

**B**

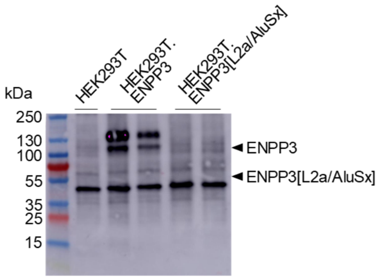


**Figure S8. Instability of the *ENPP3[L2a/AluSx]* protein product. (A)** Amino acid sequence alignment of canonical ENPP3 and the ENPP3[L2a/AluSx] isoform. Amino acid residues in red denote the sequence in ENPP3[L2a/AluSx] that replaces the end of the phosphodiesterase and all of the nuclease domain. **(B)** Western blot for ENPP3 in lysates from parental HEK293T cells and cells transduced to express ENPP3 and ENPP3[L2a/AluSx] (HEK293T.ENPP3 and HEK293T.ENPP3[L2a/AluSx], respectively, run in duplicate). Arrows show the theoretical mass of each isoform.

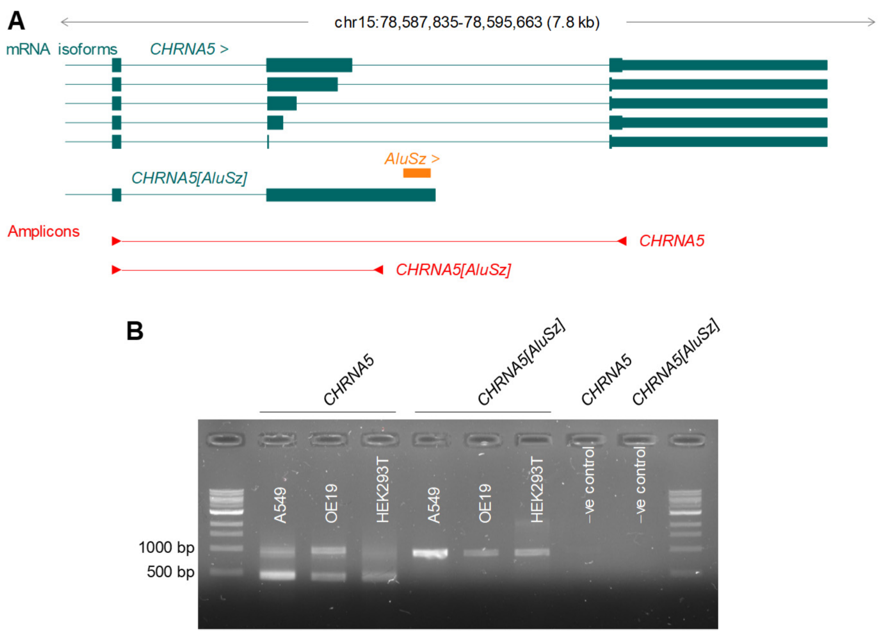


**Figure S9. PCR validation of the *CHRNA5[AluSz]* isoform. (A)** Annotated and assembled *CHRNA5* transcripts (exons 4-6 only), location of exonised *AluSz*, and amplicons used for transcript validation. **(B)** Gel electrophoresis of amplicons of *CHRNA5[AluSz]* or canonical *CHRNA5* isoforms amplified from the indicated cell lines. *CHRNA5* amplicons of different sizes correspond to canonical isoforms utilising different splice donor sites in exon 5.

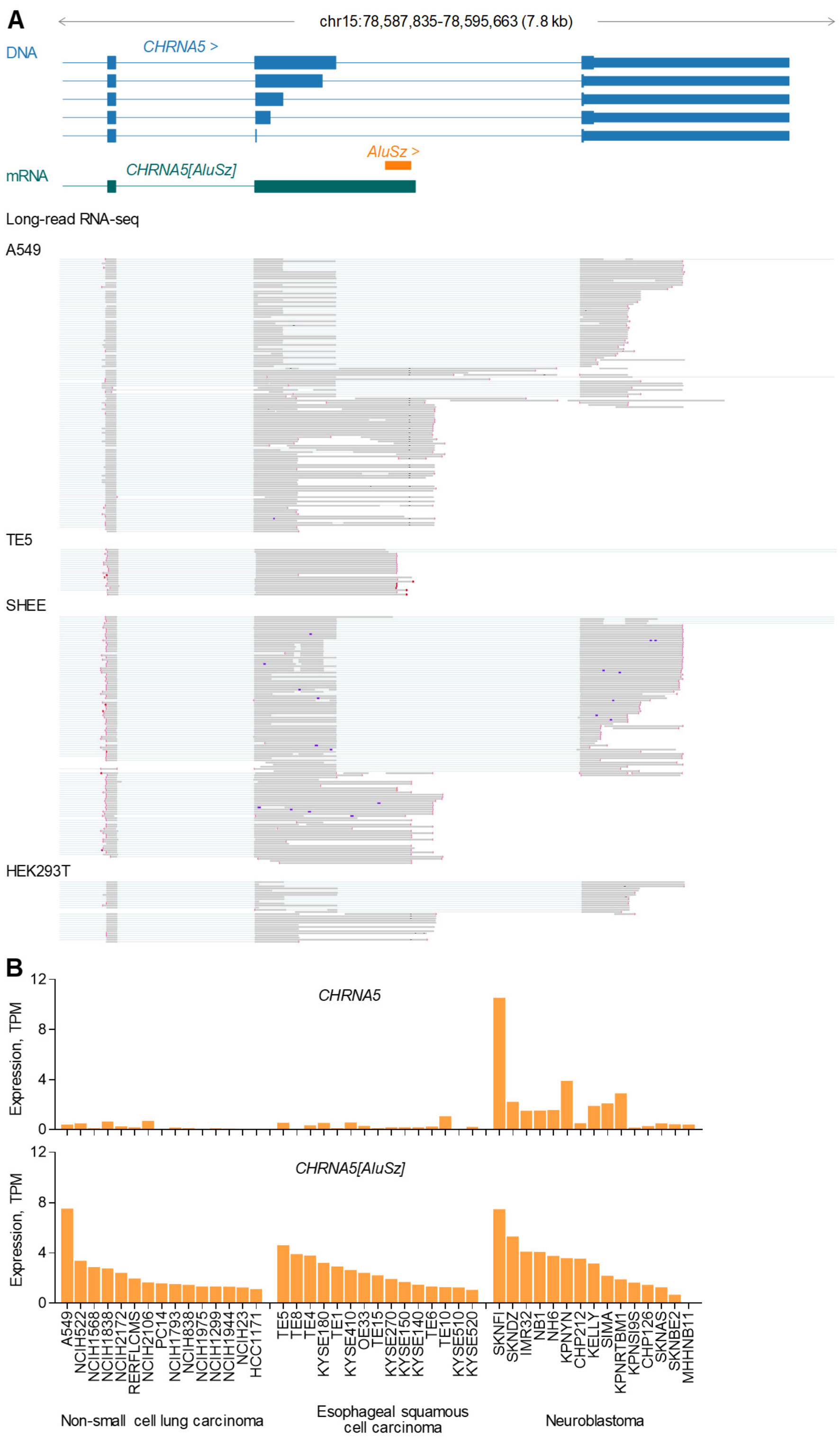


**Figure S10. Balance of *CHRNA5* isoform expression in cancer cell lines. (A)** Gene structure (exons 4-6 only), location of exonised *AluSz*, assembled *CHRNA5[AluSz]* transcript, and long-read RNA-seq data from HEK293T and A549 cells (https://github.com/GoekeLab/sg-nex-data), and esophageal squamous cell carcinoma TE5 and normal immortalized esophageal squamous epithelial SHEE cells (PRJNA515570). **(B)** *CHRNA5* and *CHRNA5[AluSz]* isoform expression (TPM) in RNA-seq data from the indicated cell lines in CCLE.

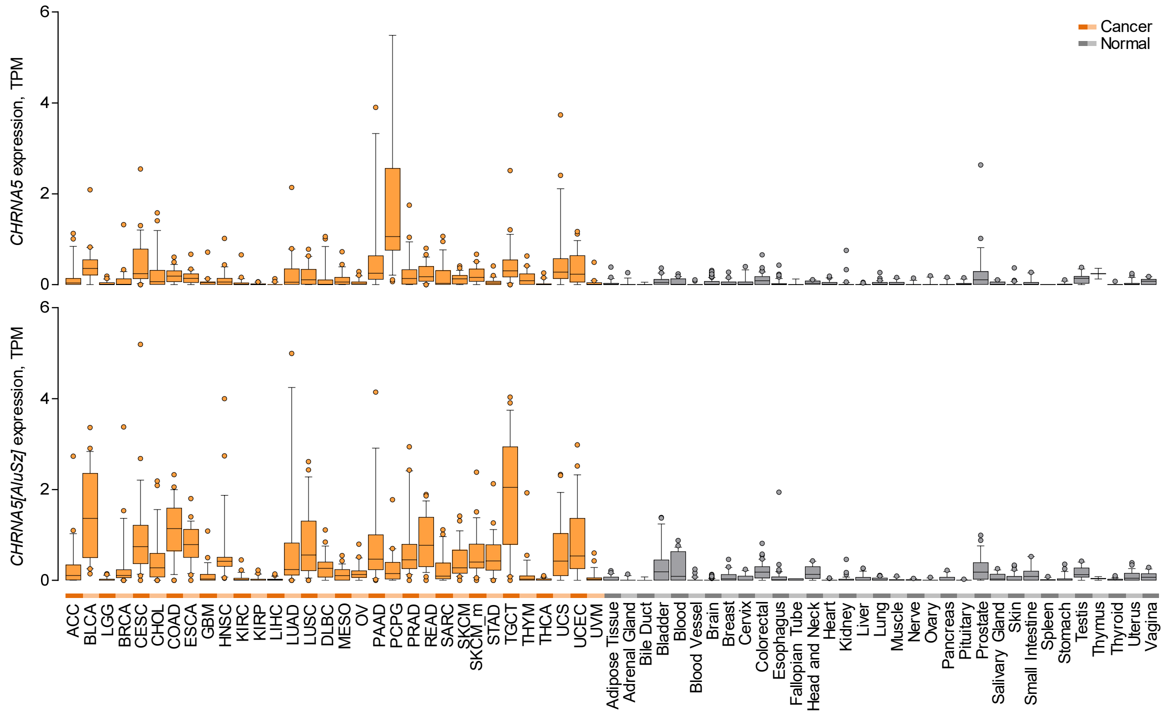


**Figure S11. *CHRNA5* and *CHRNA5[AluSz]* expression across cancers and normal tissues.** Canonical *CHRNA5* and *CHRNA5[AluSz]* isoform expression (TPM) in the indicated TCGA cancer types (n = 24 per type) and in normal tissues from GTEx (n = 2-156 per tissue type). For canonical *CHRNA5*, only the isoform with the full-length exon 5, encoding the full-length protein, is included. Box plots denote median value and quartiles, whiskers denote 1.5x the interquartile range, and individual points denote outliers.

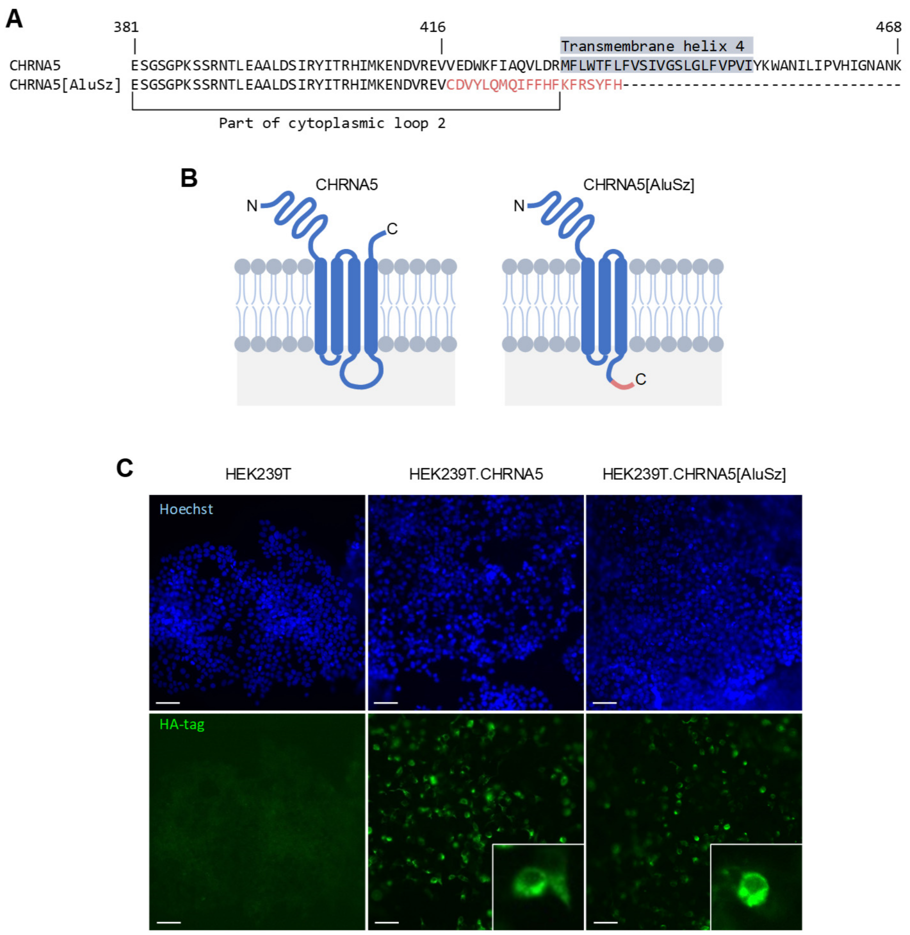


**Figure S12. Sequence and expression of the CHRNA5[AluSz] protein. (A)** Amino acid sequence alignment of canonical CHRNA5 (last 88 amino acids only) and the CHRNA5[AluSz] isoform. Amino acid residues in red denote the sequence in CHRNA5[AluSz] that replaces the end of the last transmembrane helix and extracellular C-terminal sequence of the canonical isoform. **(B)** Schematic representation of canonical CHRNA5 and the CHRNA5[AluSz] isoform. **(C)** Immunofluorescence detection of cell surface expression (in non-permeabilised cells) of HA-tagged canonical CHRNA5 and CHRNA5[AluSz] isoforms, overexpressed in HEK293T cells. Images are from a single experiment (scale bar=100µm, insets 5× magnification).

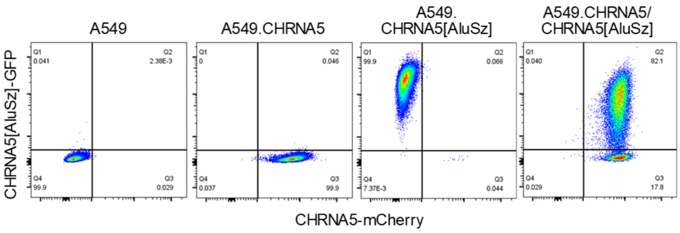


**Figure S13. Establishment of A549 cells expressing *CHRNA5* and *CHRNA5[AluSz]*.** Flow cytometric detection of fluorescent reporter expression in parental A549 cells and A549 cells expressing the canonical *CHRNA5* isoform and mCherry reporter, the *CHRNA5[AluSz]* isoform and GFP reporter or both isoforms.

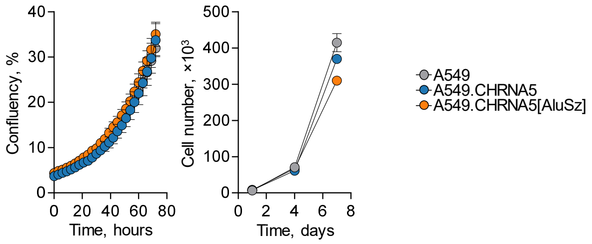


**Figure S14. Effect of *CHRNA5* and *CHRNA5[AluSz]* expression on A549 *in vitro* growth.** Mean confluency (±SEM, n=6 from 1 experiment) (*left*) and mean cell number (±SEM, n=2 from 1 experiment) (*right*) of parental A549 cell cultures and those of A549 cells expressing the canonical *CHRNA5* or the *CHRNA5[AluSz]* isoform.
